# Small-molecule allosteric activator of ubiquitin-specific protease 7 (USP7)

**DOI:** 10.1101/2025.03.14.643379

**Authors:** Isabella Jaen Maisonet, Mona Sharafi, Emilie J. Korchak, Andres Salazar-Chaparro, Ariana Bratt, Tvesha Parikh, Anthony C. Varca, Binita Shah, Michael Darnowski, Mia Chung, Wei Pin Teh, Jianwei Che, Irina Bezsonova, Sara J. Buhrlage

## Abstract

Ubiquitin-specific protease 7 (USP7) is a deubiquitylase essential for cell homeostasis, DNA repair, and regulation of both tumor suppressors and oncogenes. Inactivating USP7 mutations have been associated with Hao-Fountain Syndrome (HAFOUS), a rare neurodevelopmental disorder. Although a range of USP7 inhibitors have been developed over the last decade, in the context of HAFOUS as well as oncogene regulation, USP7 activators may represent a more relevant approach. To address this challenge, we report the discovery and characterization of a small-molecule activator of USP7 called MS-8. We showed that MS-8 activates USP7 by engaging the allosteric C-terminal binding pocket of USP7, thus mimicking the allosteric autoactivation by the USP7 C-terminal tail. We observed that MS-8 engages and activates mutant USP7 in a cellular context, impacting downstream proteins. Taken together, our study provides validation of the USP7 activator that paves the way towards novel activation-driven USP7 pharmacology.

## INTRODUCTION

Deubiquitylating enzymes (DUBs) are a family of more than 100 isopeptidases that catalyze the removal or trimming of post-translational ubiquitin tags from target proteins.^1^ DUBs balance ubiquitin signaling, thereby playing a central role in localization, protein-protein interactions, and abundance of cellular proteins.^1–3^ Ubiquitin-specific protease 7 (USP7) is a 128 kDa cysteine protease in the DUB family.^4^ USP7 has been the focus of major inhibitor development campaigns, including several from our lab, due to its well-studied role in the MDM2-p53 pathway in cancer, as well as its involvement in the regulation of tumor suppressor PTEN, Wnt/β-catenin, and NF-kB signaling pathways. ^5–7^ Recently, several roles of USP7 in neurodevelopment have been reported, including the regulation of WASH-mediated actin assembly and activation of the polycomb repressive complexes.^8–11^ Additionally, inactivating mutations in the *USP7* gene were recently identified as causal for the rare neurodevelopmental disorder, Hao-Fountain Syndrome (HAFOUS).^9,12^ Taken together, USP7 is an essential regulator of protein homeostasis, cellular signaling, growth and proliferation, and targeting USP7 may have therapeutic potential.

In cells, USP7 is maintained in an ‘inactive’ state and is activated via interactions with ubiquitinated substrates and its own 19-residue C-terminal tail that allosterically promote rearrangement of the catalytic triad into a catalysis-competent conformation.^13–16^ In general, USP7 inhibitors reported thus far, including XL-188 developed in our lab, inhibit their target by locking the enzyme in its inactive conformation. ^17^ These compounds were shown to exhibit anti-cancer effects in a wide range of cell-based and animal models of cancer, including multiple myeloma, colorectal carcinoma, bone osteosarcoma, Ewing sarcoma, breast and ovarian cancer.^5,18–21^ More recently, USP7 inhibitors served as starting points for development of proteolysis targeting chimeras (PROTACs) that, instead of inhibiting, induce degradation of USP7.^22,23^ USP7 inhibitors were also used to design DUB-targeting chimeras (DUBTACs), bifunctional molecules that recruit USP7 deubiquitylase activity to targets of interest, resulting in their deubiquitylation and stabilization.^24^ However, as USP7 is also reported to play tumor suppressive roles,^25,26^ small molecules that activate USP7 may also be of interest in that context, as well as potential therapies for HAFOUS. Furthermore, USP7 activators could serve as building blocks towards the next generation of DUBTACs with enhanced activity.

Here, we describe rigorous validation and mechanistic evaluation of MS-8, a molecule identified in a large-scale screening effort aimed at DUB inhibitor discovery, as a selective activator of USP7. Using biochemical and biophysical strategies, we demonstrated that MS-8 mimics the USP7 mechanism of allosteric autoactivation, and we confirmed that this molecule enhances the activity of HAFOUS mutant USP7 variants. We also showed that MS-8 activates USP7 in cells, resulting in down-stream stabilization of its substrates. Taken together, we provide a proof-of-concept for USP7 activation, thus highlighting the potential for developing activators for other members of the DUB family.

## RESULTS

### MS-8 activates USP7 in vitro

To identify potential USP7 activators, we reanalyzed data from our previous inhibitor-focused screen^27^, and identified several candidate selective USP7 activators. The compounds were resynthesized in-house or commercially purchased, and their potency against USP7 measured in dose-response in an in vitro fluorogenic ubiquitin-rhodamine (UbRho) assay. MS-8 (**Figure 1A**) was chosen for further study because it activated both full-length and catalytic domain USP7 in a dose-dependent manner (**Figure 1B**). As initially observed in the high-throughput screen, MS-8 was found to be selective for USP7 when tested against other DUBs (**Figure 1C, Figure S1A**). Moreover, MS-8 did not activate USP47, which shares the highest sequence homology to USP7 amongst the USP family. MS-83 activation of USP7 was orthogonally validated in two assays. First, MS-8 increased the rate of USP7 labeling with the activity-based probe, ubiquitin-propargylamine (Ub-PA), using both catalytic domain and full-length USP7 *in vitro* (**Figure 1D**). Additionally, MS-8 increased the rate of USP7 cleavage of K48-linked and K63-linked di-ubiquitin, but not of linear di-ubiquitin. This indicates that MS-8 activates the catalytic function of USP7 but not beyond its endogenous substrate scope (**Figure 1E**), as USP7 is reported to not recognize methionine-linked linear ubiquitin chains.^28^ As a result of a small medicinal chemistry campaign in our lab, we identified MS-180 as a negative control for MS-8. (**Figure S2A-F**). Chiral separation of MS-8 revealed that USP7 shows a preference for one enantiomer (P1) over the other (P2) (**Figure S1D-E**). However, given that the MS-8-P2 enantiomer maintains some USP7 activation, we chose to continue with MS-180 as our negative control, to give us a larger degree of distinction.

**Figure 1.**
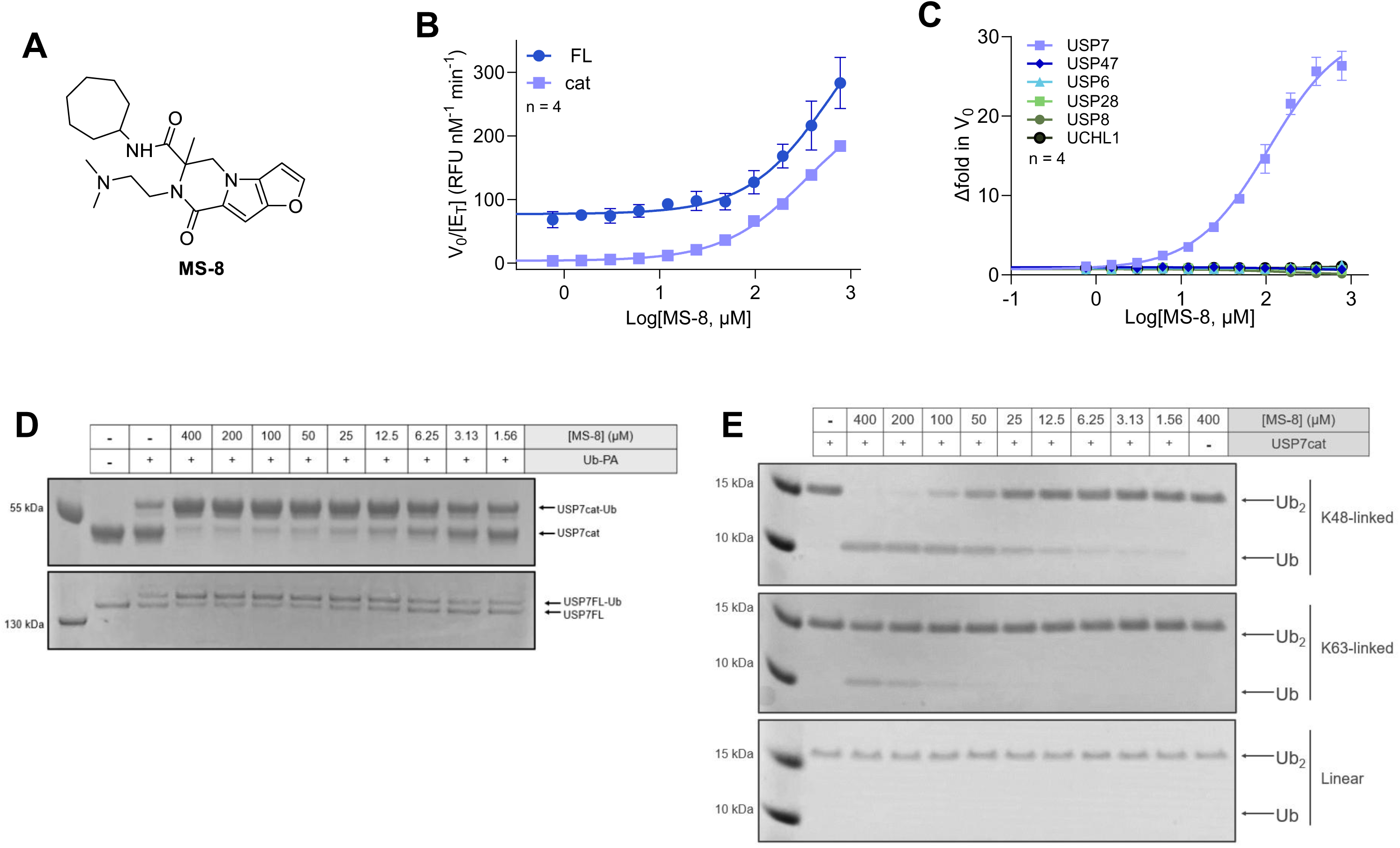
MS-8 is a selective USP7 activator. (**A**) Chemical structure of MS-8. (**B**) USP7FL and catalytic domain initial velocities when supplemented with MS-8 in dose response. Activity of DUBs assessed using ubiquitin-rhodamine assay. n = 4. (**C**) MS-8 dose response DUB selectivity assay using fluorogenic ubiquitin-rhodamine as substrate. n = 4. (**D**) Covalent labeling of USP7 catalytic domain and full length with ubiquitin-propargylamine after 2 hours, with and without MS-8 dosing. (**E**) Di-ubiquitin cleavage with USP7 catalytic domain and MS-8 dosing after 2 hours. K48-, K63- and methionine-linked di-ubiquitin were each assessed.

### MS-8 binds the allosteric C-terminal peptide binding pocket on USP7

We paired *in silico* computer modeling with NMR chemical shift perturbation mapping to identify the binding pocket for MS-8. Using the crystal structures of USP7 bound to ligands, we constructed models of the USP7 catalytic domain for molecular docking and molecular dynamic (MD) simulations. To start, we docked MS-8 into three of USP7’s best characterized pockets (**Figure 2A**). Two of the pockets, the distal and S3-S5, have been previously shown to bind small-molecule inhibitors (PDB: 5UQX and PDB: 5VS6, respectively).^5,29^ We additionally conducted docking of MS-8 into the reported allosteric pocket (inactive form of USP7, using PDB: 5UQX) where the C-terminal tail of USP7 binds for autoactivation, herein called the C-terminal peptide binding (CPB) pocket. Docking models demonstrated that MS-8 fit well within the CPB pocket and distal pockets, and not the S3-S5 pocket (**Table S1A**).

**Figure 2.**
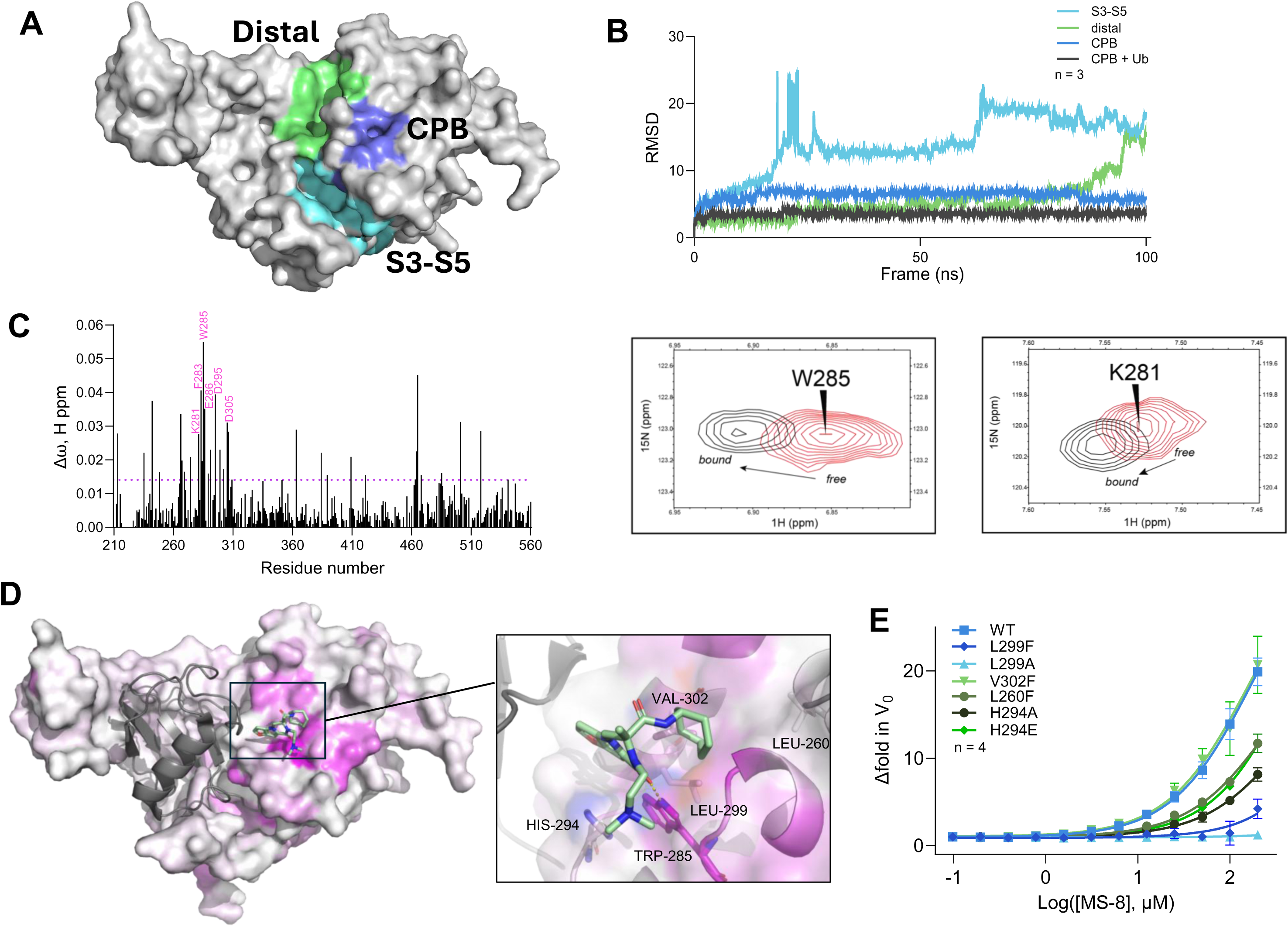
MS-8 binds allosteric CPB pocket. (**A**) USP7 catalytic domain (PDB: 5JTV) with previously described pockets colored; distal ubiquitin-binding pocket (green), S3-S5 catalytic cleft (light blue), and C-terminal peptide binding (CPB) pocket (dark blue). (**B**) RMSD of MS-8 throughout 100 ns molecular dynamic simulation, starting from top ranked docking poses in each pocket. Data shows mean RSMD of three separate simulations (n = 3). (**C**) Bar graph depicting per-residue NMR chemical shift perturbations (Δω) in the ^15^N-TROSY spectrum of the USP7 catalytic domain upon the addition of a 4-fold molar excess of MS-8. Overlaid spectra for the free (red) and MS-8-bound (black) states are shown for the W285 and K281 amide peaks as examples. The complete spectrum overlay is provided in Supplemental Figure S4A. (**D**) Overlay of MS-8 in CPB pocket from MD simulation, with NMR chemical shift perturbation data from Figure 2C mapped onto USP7 catalytic domain structure (PDB: 4M5W). The pinker, the larger the perturbations. (**E**) Dose-response of MS-8 against USP7 catalytic domain with mutations in CPB pocket assessed by ubiquitin-rhodamine assay. n = 4.

To further evaluate the binding, the top ranked docking poses in each pocket were selected for post-docking analysis and subsequent MD simulations, in triplicates (**Figure 2A-B**). Throughout the MD simulation time (100 ns), MS-8 was stable in the CPB pocket, with an average RMSD score of ∼ 6 Å and MMGBSA of -35 kcal/mol. Further, MS-8 was unstable and dissociated from both the distal and S3-S5 pockets during the MD simulation time. Interestingly, the thorough analysis of the trajectories from the last 30 ns MD revealed that after MS-8 dissociates from both pockets, it eventually anchors on to the CPB pocket (**Supplemental Video 1**), in three replicates. As a control, we additionally performed docking and MD simulation with the reported distal pocket inhibitor, GNE-6776,^29^ which maintained its crucial interactions in distal pocket over the course of MD simulation.

Considering the conformational difference between the active and inactive state of USP7, we next investigated how the presence of ubiquitin can influence the stability of the activator binding computationally. To reach this goal, we ran an additional MD simulation in the presence of ubiquitin (PDB: 5JTV) and observed significant improvements in MS-8/USP7 complex stability in the CPB pocket compared to inactive complex with average RMSD score of ∼3 Å and MMGBSA of -47 kcal/mol (**Figure 2B, Supplemental Video 2**).

The model indicates that MS-8 engages in hydrophobic interactions of the cycloheptyl ring with V302, L299, and F283, which anchors MS-8 to the protein, and a hydrogen-bond between a MS-8 carbonyl and W285. We confirmed the *in silico* results with ^15^N/^2^H isotope labeled protein nuclear magnetic resonance (NMR) spectroscopy (**Figure 2C, Figure S4A-B**). The residues with the largest chemical shift perturbations in the presence of MS-8 (W285, F283, and K281) belong to the same pocket we identified in our computational studies (**Figure 2D**). Subsequent point-mutagenesis of key residues in that pocket decreased (L260F, H294A/E) or fully abrogated (L299A, L299F) MS-8-mediated activation of USP7 catalytic domain, except for V302F, which had no effect (**Figure 2E**). This could indicate that V302 of USP7 is only primarily involved in contributing to the hydrophobicity of the pocket which is not sufficiently perturbed by a V to F mutation. Taken together, our docking, MD simulations, protein NMR, and pocket mutagenesis studies support our conclusion that MS-8 binds the allosteric CPB pocket on USP7.

### MS-8 competes with the C-terminal tail of USP7 for binding in the CPB pocket

Given that MS-8 occupies the same pocket as the USP7 C-terminal tail in the autoactivated conformation, we hypothesized that MS-8 activates USP7 in a similar manner as the C-terminal tail. We superimposed our model of MS-8 bound to USP7 with the previously reported X-ray crystal structure that features the C-terminal tail peptide bound to the CPB pocket of USP7 (PDB: 5JTV).^16^ The overlay revealed several chemical parallels between MS-8 and the tail (**Figure 3A**). Specifically, the dimethylamine on MS-8 overlays with K1099 on the C-terminal tail, and the cycloheptyl ring makes similar hydrophobic interactions with the pocket as I1100. Additionally, the backbone carbonyl of K1099 aligns with the carbonyl on MS-8, both of which form hydrogen bonds with W285. Previous work showed that mutations at these residues (K1099 and I1100) each resulted in a greater than 90% reduction in USP7 activity.^16^ These parallels suggest that MS-8 and the C-terminal tail compete for binding.

**Figure 3.**
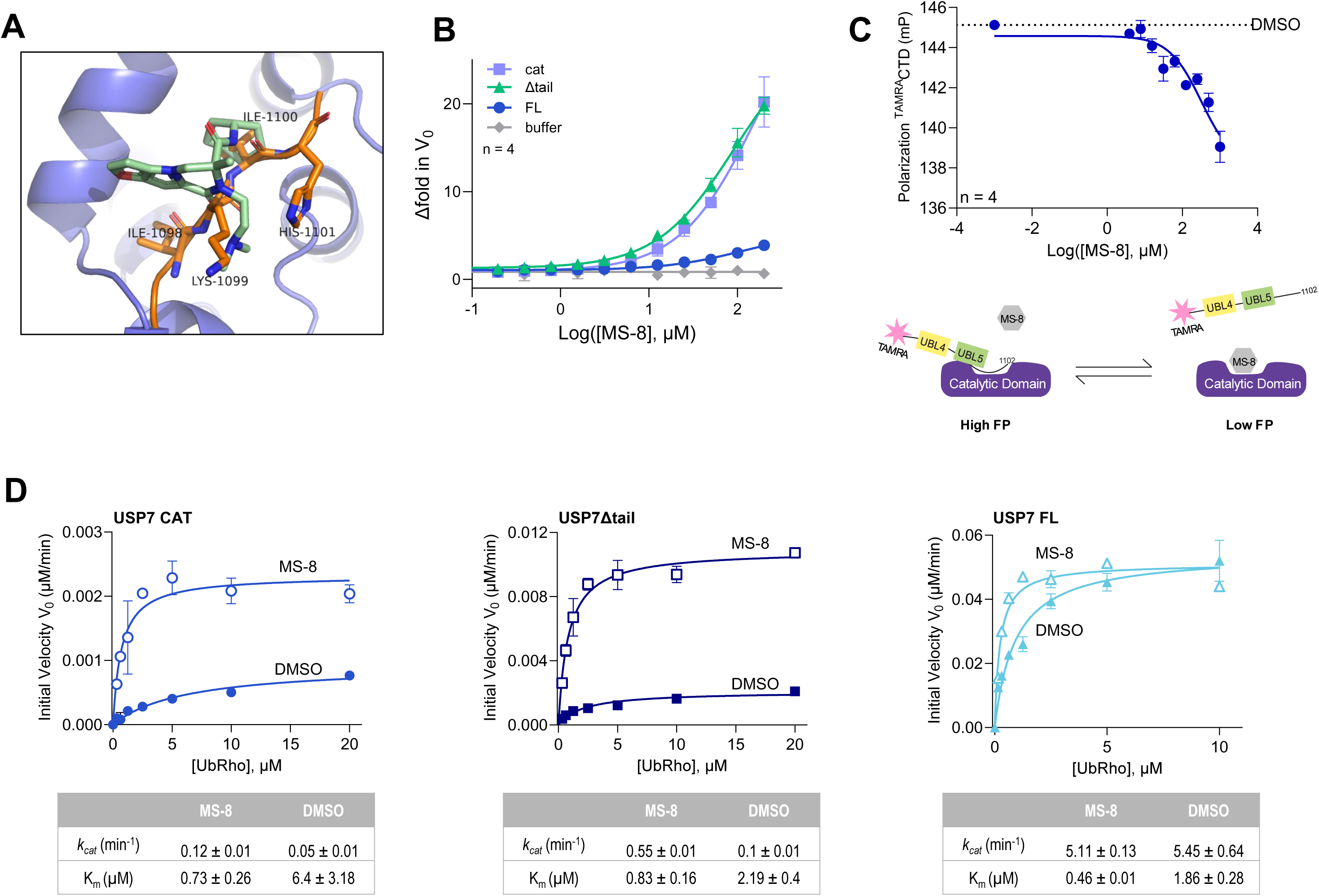
MS-8 competes with the C-terminal tail for binding. (**A**) Overlay of MS-8 (green) from MD simulations in USP7 CPB pocket (dark blue) with C-terminal tail peptide (orange) from PDB: 5JTV (**B**) MS-8 ubiquitin-rhodamine assay with USP7 catalytic domain, full length, and ΔC-terminal tail. n = 4. (**C**) Competition fluorescence polarization assay between TAMRA-labeled USP7 C-terminal domains (CTD, aa 883-1102) and MS-8 for binding the USP7 catalytic domain. Binding of TAMRA-USP7-CTD to USP7 catalytic domain results in high FP. n = 4. (**D**) Michaelis-Menten kinetic assessment of USP7 constructs with ubiquitin-rhodamine as substrate, with either DMSO or MS-8 (31.3 µM). Full dose-response and table for each USP7 construct in Figure S3A-C.

To confirm this, we first generated a USP7Δtail mutant and observed that MS-8 activates this construct to the same extent as the USP7 catalytic domain (**Figure 3B**). We also examined the effect of MS-8 on the kinetics of reactions catalyzed by USP7 catalytic domain, USP7 full-length, and the USP7Δtail construct (**Figure 3D, Figure S3A-C**). For both the USP7 catalytic domain and USP7Δtail, we observed a decrease in *K_m_* and a significant increase in *V_max_*, while the full length USP7 had a more subtle decrease in *K_m_* with MS-8 at low concentrations. Further, we measured the direct competition between MS-8 and the C-terminal tail via a competitive fluorescence polarization (FP) assay using the C-terminal domains (CTD) of USP7 (residues 883-end) N-terminally labeled with 5-TAMRA. The TAMRA-USP7_CTD_ probe binding to the USP7 catalytic domain results in a high FP signal. After dosing MS-8, we measured a decrease in FP, as the small molecule competes off the TAMRA-USP7_CTD_ (**Figure 3C**). Taken together, these results show that MS-8 engages the same binding site as the C-terminal peptide, the CPB pocket, and uses similar molecular features to engage with the same contact points as the C-terminal peptide. As seen in our FP studies, MS-8 is able to outcompete the C-terminal peptide. It is then reasonable to propose that MS-8 induces comparable allosteric changes as the C-terminal tail, ultimately activating USP7 deubiquitylase activity.

### MS-8 mimics USP7 mechanism of allosteric autoactivation

To test this possibility further, we used TAMRA-labeled ubiquitin and measured the binding affinity of ubiquitin to USP7 constructs using FP. We observed little to no change in the binding affinity of ubiquitin in the presence of MS-8, concluding that MS-8 does not activate USP7 by affecting its affinity for substrate (**Figure 4A, Table S2C**). This aligns with the mechanism of the C-terminal tail, which also does not affect the binding affinity of USP7 for ubiquitin.^14^

**Figure 4.**
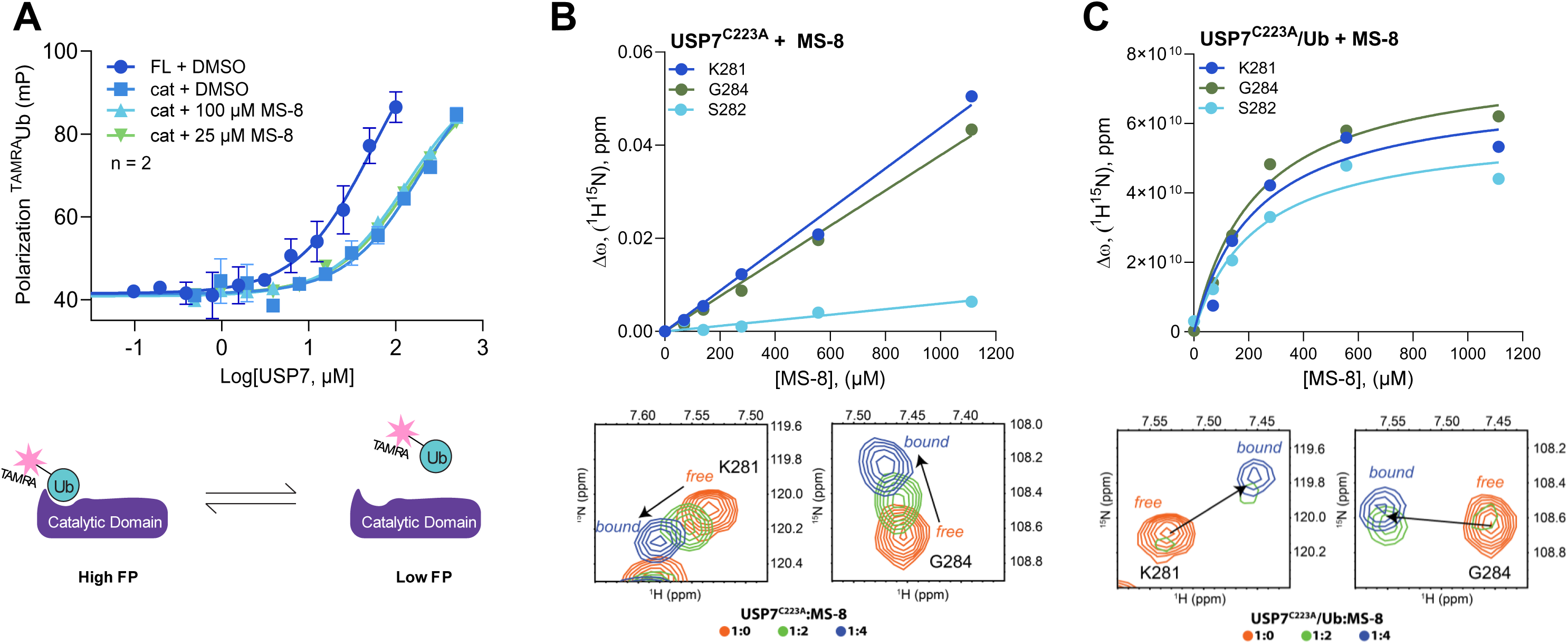
MS-8 mimics USP7 C-terminal tail mechanism of activation. (**A**) Fluorescence polarization of TAMRA-labeled ubiquitin with USP7 full length, catalytic domain, or catalytic domain and MS-8 at 100 µM or 25 µM. n = 2. (**B-C**) NMR titration of free and ubiquitin-bound USP7-C223A with MS-8. Representative chemical shift perturbations (Δω) in the ^15^N-TROSY spectrum of the catalytic domain are plotted as a function of MS-8 concentration. Spectral changes during the titration process are shown below each plot as illustrative examples.

Furthermore, the C-terminal tail of USP7 was previously reported to bind sequentially following substrate binding and activate the DUB using a mechanism similar to a classic “induced fit”.^14^ To test whether MS-8 employs a similar activation mechanism, we used NMR chemical shift perturbation assays to monitor the binding of MS-8 to either the catalytic domain of USP7 or its complex with ubiquitin. These assays used the catalytic cysteine-to-alanine mutant of USP7 previously shown to stabilize other DUB/ubiquitin complexes.^30^ The changes in the ^15^N-^1^H TROSY spectrum of the C223A catalytic domain induced by the addition of MS-8 affected residues in the CPB pocket such as W285, D295, and D305 (**Figure S4C**). However, the dependence of chemical shift perturbations on the MS-8 concentration showed that the interaction is weak (**Figure 4B**). In contrast, the addition of MS-8 to the ^15^N-labeled catalytic domain loaded with ubiquitin resulted in significantly tighter binding (**Figure 4C**). These results suggest that similar to the C-terminal tail, MS-8 activates USP7 by preferentially binding to the allosteric pocket of the ubiquitin-loaded catalytic domain (**Figure 4B-E**). This model is consistent with our computational observation that ubiquitin improves the CPB pocket’s stability (**Figure 2A**).

Taken together, our data demonstrate that the mechanism of action of MS-8 mimics the endogenous autoactivation of USP7 by its C-terminal tail.

### MS-8 activates cellular mutant USP7

We next sought to determine if MS-8 could activate wildtype and mutant USP7 in a cellular context. We first assessed its cellular toxicity in both HCT116 wildtype (USP7^+/+^) and HCT116 USP7 knockout (USP7^-/-^) lines, along with the negative control MS-180. After a 24-hour incubation period, we observed no changes in viability up to a concentration of 100 µM, confirming their utility in cellular context as it allows for specific and interpretable biological experiments without confounding effects from cell death or stress responses. (**Figure S5A-B**).

Next, we examined the impact of small molecule treatment on reported USP7-regulated proteins, TRIM27 and p21, in the HCT116 USP7^+/+^ and HCT116 USP7^-/-^ cells (**Figure 5A**). As expected, both TRIM27 and p21 were impacted by the knockout of USP7. Additionally, we observed a decrease in TRIM27 and an increase in p21 upon USP7 inhibitor (XL177A) treatment in the HCT116 USP7^+/+^, but not in the cells lacking USP7. Little to no difference in TRIM27 or p21 levels were observed upon treatment with MS-8 or MS-180, consistent with the limited activation seen *in vitro* for USP7 wild-type (**Figure 1B, S1B-C**). This is most likely due to the fact that USP7 wild type is already a highly active enzyme resulting in a narrow window of further activation by MS-8. Given these results, we transiently transfected USP7 WT and USP7Δtail into the HCT116 USP7^-/-^ line to confirm MS-8-mediated activation as observed *in vitro*. Markedly, only in the USP7Δtail transfected cells did MS-8 treatment results in a decrease in p21, a downstream result of p53 decrease by USP7-mediated rescue of MDM2 (**Figure 5C**).

**Figure 5.**
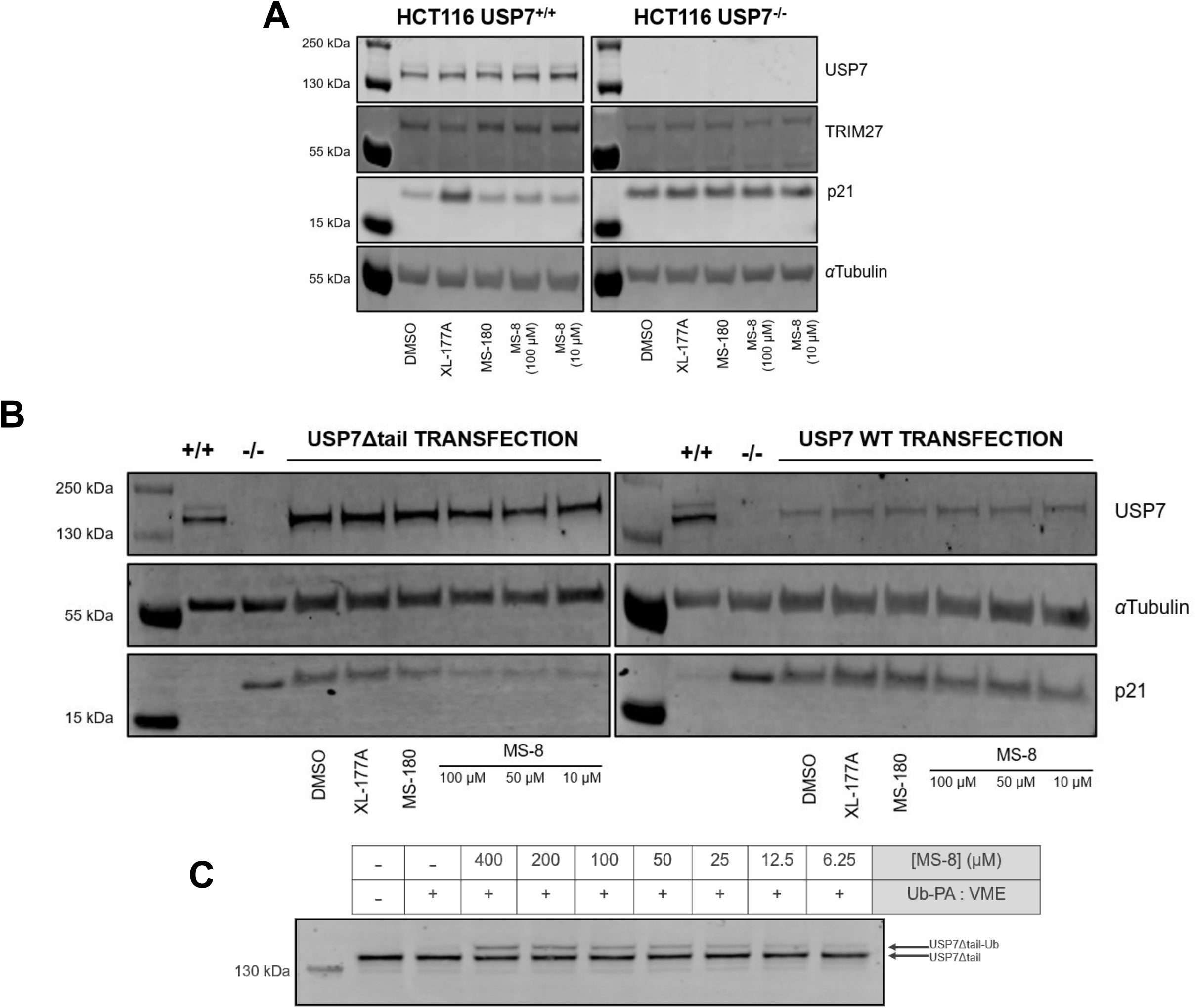
MS-8 activates cellular mutant USP7. (**A**) Western blot analysis of USP7, TRIM27, and p21 levels from HCT116 USP7^+/+^ cells and HCT116 USP7^-/-^ cells. Cells were treated with either DMSO, XL-177A (10 µM), MS-180 (100 µM), or MS-83 (100 µM or 10 µM) for 24 hours. (**B**) HCT116 USP7 parental (+/+), HCT116 USP7 knockout (-/-) and USP7 knockout transfected with either USP7Δtail or USP7 WT. Cells were treated with either DMSO, XL-177A (10 µM), MS-180 (100 µM), or MS-83 (100 µM, 50 µM, or 10 µM) for 24 hours. (**C**) Covalent labeling of USP7Δtail from HCT116 USP7^-/-^ transfected with USP7Δtail lysate. Covalent labeling with 1:1 mixture of ubiquitin-propargylamine (Ub-PA): ubiquitin-vinylmethylester (Ub-VME) for 2 hours at room temperature.

We confirmed USP7Δtail target engagement by MS-8, by treating the lysate with a covalent ubiquitin-probe (**Figure 5D**). We saw MS-8 dose responsive increase in mutant USP7 labeling with the activity-based covalent probe, aligning with our *in vitro* results with this construct (**Figure 3B, 3D, S3B**). Taken together, MS-8 is able to engage and activate mutant USP7 in cells, resulting in the stabilization of a known substrate of USP7.

## DISCUSSION

Traditionally, target-based drug discovery has focused on developing inhibitors of enzymatic activity, primarily with the goal of preventing aberrant cell signaling events from taking place. However, in some cases, enzyme activation through small-molecule mediation may offer a more appropriate strategy for addressing different disease states.^31,32^ Although small molecule activators of AMP-activated protein kinases (AMPK) were described more than a decade ago,^32–34^ pharmacological activation remains in its infancy. Since then, small molecule activators have been disclosed for only a small handful of targets, many of which are kinases (e.g. glucokinase (GK),^35^ cGMP-dependent protein kinase (PKG),^36^ lysophospholipase-like 1 (LYPLAL1),^37^ and phosphoinsositol 3-kinase α (PI3Kα)^38^). Small molecule activators are rare in part due to the difficulty of separating bona fide activators from false positives in high-throughput screens, and the lack of defined methods for validating activation.

These reported enzymes amenable to small molecule-mediated activation all share the presence of an allosteric pocket that, when engaged with an endogenous or exogenous activator, induces conformational perturbations that translate into increased enzyme activity. In our example, the crucial conformational rearrangement that results in the activation of USP7 involves the movement of the catalytic triad from an auto-inhibited state to an active arrangement. Endogenously, USP7’s C-terminal tail binds to the allosteric pocket near the active site to regulate USP7 activity.^13,14,16^ We now describe MS-8, a small-molecule activator that increases USP7 catalytic activity by binding to the same allosteric C-terminal peptide binding (CPB) pocket, and, thus, mimicking the mechanism of the C-terminal dependent autoactivation.

Initially identified from a high-throughput screen, MS-8 was confirmed as a bona fide USP7 activator after rigorous validation in various USP7 activity assays. MS-8 activates both USP7 catalytic domain and full-length selectively when tested against a panel of other DUBs, including its closest homolog, USP47. We provide evidence that MS-8 binds the allosteric CPB pocket of USP7, driven by hydrophobic interactions and a hydrogen bond between MS-8 and W285. Additionally, we demonstrate that this small molecule competes with the C-terminal tail for binding. Mechanistically, we confirmed that MS-8 does not activate by improving the binding affinity of ubiquitin to USP7, and that MS-8 binds preferably to the ‘active’ ubiquitin-bound USP7, aligning with USP7 literature for the C-terminal tail. Beyond an *in vitro* context, we demonstrate MS-8’s ability to activate the catalytically inactive USP7Δtail construct in cells, as evidenced by the dose-response increase in Ub-PA labeling upon treatment with MS-8, and the decrease in p21 levels, a direct transcriptional target of p53 in a well-annotated pathway of USP7 activity. During the preparation of this manuscript, two FDA-approved drugs, Sertraline (Zoloft®), an antidepressant that potently targets the serotonin reuptake pathway, and Astemizole, an antihistamine drug, were proposed to, in addition to their well-accepted mechanism of action via the serotonin/5-HT transporter and the histamine H1 receptor, respectively, also act as activators of USP7.^39^ Similar to MS-8, these two compounds also act by engaging the CPB pocket. Therefore, the allosteric CPB pocket of USP7 appears to represent the hotspot for modulation of USP7 activity.

MS-8 offers a starting point for developing next-generation activators of USP7. Although USP7 is primarily viewed from the perspective of inhibitor development (see XL-177A, FT671, and ALM5 for examples of well-validated USP7 inhibitors),^6,18,19^ USP7 activation represents an important strategy for understanding *USP7* haploinsufficiency in HAFOUS and treating those diseases that are associated with *USP7* loss-of-function mutations. Furthermore, several other DUBs, like BAP1 and OTUD1, are reported to have loss-of-function pathogenic mutations that could benefit from the development of small molecule activators.^40,41^ Our work here offers a framework for identifying and validating DUB small-molecule activators, which could greatly impact how we approach probing DUB biology, both in the context of wild-type and loss-of-function pathologies.

Additionally, DUB activators may have important implications for developing a new pharmacological modality for targeted protein stabilization (TPS). Thus far, TPS strategies have employed DUBTACs (DUB TArgeting Chimeras), heterobifunctional small molecules that consist of a DUB binder linked to a ligand for the protein target of interest, to stabilize the levels of the target by deubiquitylating it and preventing its proteasomal degradation. ^42^ One of the key design principles for these novel modalities, that has been noted in the proof-of-concept studies, is that DUB-recruiting ligands should bind to allosteric sites, in order to leave access to the active site unobstructed. Importantly, for these agents to work, DUB binding ligands should either be neutral (not inhibit) or enhance DUB activity. From these two standpoints, MS-8 represents a promising starting point for developing USP7-directed TPS strategies. Overall, this work shows that targeting the conformational mechanisms of DUBs could be the key to accessing this therapeutic target family and their pathways.

## ACKNOWLEDGMENTS

We would like to acknowledge generous support from the Helen Gurley Brown Foundation, the Jean Strouse Sharf and Lisa Sharf Green Cancer Research Fund, the Foundation for USP7-Related Diseases Grant, NIH R35GM128864, and NIH 1R21NS135343. We would like to thank Dr. Milka Kostic for the valuable discussions of the manuscript. We thank the MRC PPU Reagents and Services facility (MRC I PPU, College of Life Sciences, University of Dundee, Scotland, mrcppureagents.dundee.ac.uk) for the pGEX-6-USP7-883-end plasmid.

## AUTHOR CONTRIBUTIONS

I.J.M. and S.J.B wrote the manuscript and prepared figures. I.J.M., M.S., S.J.B., I.B. conceptualized the project. I.J.M. designed and performed biochemical and kinetic assays. M.S. designed and performed computational studies. E.J.K. and T.P. designed and performed NMR chemical shift perturbation assays. A.S.C. designed and performed cellular testing. A.C.V. identified initial hit from screening data. I.J.M., A.S.B., E.J.K., B.S., and A.S.C. designed and generated USP7 mutants. I.J.M., A.S.B., E.J.K, and B.S. expressed and purified protein. M.S. designed synthetic routes for MS-8 and MS-180. M.S., W.P.T., and M.D. performed chemical synthesis. M.C. evaluated chiral compounds. J.C. provided computational software licenses and valuable training. S.J.B. and I.B. acquired funding, reviewed and edited the manuscript. All authors read and approved the final version prior to submission.

## DECLARATION OF INTERESTS

S.J.B is a founder, SAB member, and equity holder of Entact Bio and receives or has received sponsored research funding from Novartis Institutes for BioMedical Research, AbbVie, Kinogen, TUO Therapeutics, Takeda, and Pivotal Life Sciences

I.J.M., M.S., S.J.B, and I.B. are named inventors on patent related to this work.

## STAR METHODS

| REAGENT or RESOURCES | SOURCE | IDENTIFIER |
| --- | --- | --- |
| AnaTag™ 5-TAMRA Protein Labeling Kit | AnaSpec | Cat. AS-72063, Lot. 1029 |
| HAUSP (D17C6) XP Rabbit mAb | Cell Signaling | Cat. 4833, Lot. 1 |
| p21 Waf1/Cip1 (12D1) Rabbit mAb | Cell Signaling | Cat. 2947, Lot. 12 |
| TRIM27 Rabbit PolyAb | Proteintech | Cat. 12205-1-AP, Lot. 00085906 |
| beta-Actin Rabbit Ab | Cell Signaling | Cat. 4967, Lot. 15 |
| Anti-alpha tubulin | Sigma-Aldrich | Cat. T5168, Lot. 0000286919 |
| IRDye® 800CW Goat anti-rabbit | Li-Cor | Cat. 926-32211, Lot. C80118-05 |
| IRDye® 680LT Goat anti-mouse | Li-Cor | Cat. 926-68020, Lot. D30418-15 |

| Bacterial and virus strains |  |  |
| --- | --- | --- |
| NEB® 10-beta Competent *E. coli* (High Efficiency) | New England BioLabs Inc | Cat. C3019H |
| BL21(DE3) Competent *E. coli* | Invitrogen | Cat. C600003 |
| **Chemicals, peptides, and recombinant proteins** |  |  |
| Ubiquitin-Rhodamine 110 (Ub-Rho) | R&D Systems | Cat. U-555-050, Lot. DBFH2723111 |
| Ubiquitin-propargylamine (Ub-PA) | UbiQ | Cat. UbiQ-076 |
| Ubiquitin-vinylmethylester (Ub-VME) | UbiQ | Cat. UbiQ-054 |
| USP6cat-GST (529 – 1406) | Ubiquigent | Cat. UBI-64-0045-050 |
| Ubiquitin, TAMRA-labeled | LifeSensors, Inc. | Cat. SI-270T, Lot. SI-48431.001 |
| K48-linked (Ub<sub>2</sub>) di-ubiquitin | LifeSensors, Inc. | Cat. SI-4802, Lot. SI-48526.002 |
| K63-linked (Ub<sub>2</sub>) di-ubiquitin | LifeSensors, Inc. | Cat. SI-6302, Lot. SI-48468.001 |
| Di-ubiquitin (linear) GST-cleaved | Ubiquigent | Cat. 60-0115-050, Lot. 30108 |
| **Experimental models: cell lines** |  |  |
| HCT116 | ATCC® | CCL-247™ |
| HCT 116 USP7 (-/-) | Horizon Discovery | HD R02-028 |
| **Recombinant DNA** |  |  |
| pDARMO-USP7 (Flag-tagged, aa 1-1102) | This paper | N/A |
| pDARMO-USP7Δtail (Flag-tagged, aa 1-1083) | This paper | N/A |
| pCI-neo-USP7 (Flag-tagged, aa 1-1102) | Lamberto et al<sup>5</sup> | N/A |
| pCI-neo-USP7Δtail (Flag-tagged, aa 1-1083) | This paper | N/A |
| pET28aLIC-USP7 (His-tagged, aa 207-506) | This paper | N/A |
| pET28pp-USP47 (His-tagged, aa 182-603) | This paper | N/A |
| pETDuet-USP28 (aa 149-703) | This paper | N/A |
| pET28aLIC-USP8 (His-tagged, aa 742-1110) | Varca et al<sup>27</sup> | N/A |
| pGEX6P1-UCHL1 (aa 1-223) | Varca et al<sup>27</sup> | N/A |
| pGEX-6-USP7 (aa 883-1102) | MRC PPU Reagents and Services facility (MRC PPU, College of Life Sciences, University of Dundee, Scotland, mrcppureagents.dundee.ac.uk) | DU13957, GenBank NP_003461.2 |
| pET15b-Ubiquitin (His-tagged, a 1-76) | Pozhidaeva et al<sup>43</sup> | N/A |

| Software and algorithms |  |  |
| --- | --- | --- |
| Prism 10.3 | GraphPad Software, Inc. | https://www.graphpad.com/scientific-software/prism/ |
| Maestro | Schrodinger, Inc. | https://www.schrodinger.com/platform/products/maestro/ |
| Desmond | Schrodinger, Inc. | https://www.schrodinger.com/platform/products/ms-desmond/ |
| PyMOL | Schrodinger, Inc. | https://pymol.org/ |
| Image Studio Lite Ver 5.2 | Li-Cor Bio | https://www.licorbio.com/image-studio |
| SPARKY 3 | Goddard and Kneller et al.<sup>44</sup> | https://www.cgl.ucsf.edu/home/sparky/. |
| NMRPipe | Delaglio et al.<sup>45</sup> | https://spin.niddk.nih.gov/bax/software/NMRPipe/ |

### USP7 cloning and point mutagenesis

USP7 catalytic domain (residue 207-560) was cloned into pET28a-LIC (Structural Genomics Consortium), downstream of the N-terminal 6xHis tag and thrombin cleavage site. Point mutations were introduced to the USP7 catalytic domain construct using Q5 Site-Directed Mutagenesis Kit (New England Biolabs), with primers (**Table S2A**) obtained from Integrated DNA Technologies. PCR products were transformed into *E. coli* 10-β and mutant plasmids isolated. Mutations were confirmed by whole-plasmid sequencing (Plasmidsaurus), prior to transformation into *E. coli* BL21(DE3) for protein expression and purification.

The USP7Δtail construct (residue 1-1083) was prepared by PCR cloning using the pCI-neo Flag HAUSP vector (Addgene # 16655) as a template. Briefly, 100 ng of pCI-neo Flag HAUSP template were mixed with 10 µM of forward and reverse primers (**Table S2B**) and 2x Q5 High-Fidelity Master Mix (New England BioLabs®. # E0555S) and amplified following standard thermocycling conditions. PCR product was purified using the Monarch® DNA Gel Extraction Kit (New England BioLabs®. # T1020), transformed into NEB® 5-alpha Competent E. coli (New England BioLabs®. # C2987H), and plated on LB plates with ampicillin for antibiotic selection. USP7 full length and USP7Δtail were cloned into a pDARMO (pDarmo.CMVT_v1 was a gift from David Sabatini) expression plasmid with an N-terminal FLAG tag, for protein purification. Whole-plasmid sequencing was performed by Plasmidsaurus using Oxford Nanopore Technology with custom analysis and annotation.

USP47 catalytic domain plasmid was cloned into pET28pp from pDEST26-USP47 (Addgene# 64326). pETDuet SUMO-USP28 catalytic domain plasmid was synthesized by GeneWiz, and the plasmid for SUMO protease ULP-1 was obtained from Addgene.

### Protein purification

Catalytic domain of USP7 (wild-type and mutants), USP47, USP8, USP6, USP28, C-terminal domains of USP7 (883-end), and full-length UCHL1 were expressed in *E. coli* BL21(DE3) in LB media with antibiotic. Bacteria were grown at 37 °C in a shaker incubator and induced at an OD of 0.6-0.8 with 1 mM Isopropyl β-D-1-thiogalactopyranoside (IPTG), then incubated at 16 °C overnight. Bacteria were harvested (4 °C, 4,000 rpm, 20 min). Pellets were lysed in lysis buffer (25 mM Tris pH 8.0, 1 M NaCl, 10 mM β-mercaptoethanol (BME), and protease inhibitor cocktail (Roche)) and sonicated (3:30, 15 sec on, 30 sec off). Lysates were centrifuged (4 °C, 18,000 rpm, 20 min). Lysis supernatants were transferred to Ni-NTA beads (Qiagen) in column, incubated for 1 hour at 4 °C, then washed with lysis buffer with 25 mM imidazole. Protein was eluted with elution buffer (25 mM Tris pH 8.0, 200 mM NaCl, 300 mM imidazole, and 10 mM BME). The protein elution was concentrated to 500 µL, then further purified via FPLC on a Superdex 200 (GE Healthcare) in buffer (25 mM HEPES pH 7.5, 200 mM NaCl, and 1 mM dithiothreitol (DTT)). Protein fractions were pooled, concentrated, aliquoted and stored in -80°C.

NMR samples of the USP7 catalytic domain (residues 207-560) and its C223A mutant were ^15^N-labeled. The WT USP7 sample used for mapping the MS-8 binding site was ^15^N/^2^H-labeled. Proteins were expressed in *E. coli* BL21(DE3) cells grown in M9 minimal media with ^15^NH_4_Cl as the sole nitrogen source. Deuterated media was used for ^15^N/^2^H-labeled samples. Cells were grown at 37 °C until OD_600_ reached 0.8-1.0 o.u. Protein expression was induced with 1 mM IPTG at 20°C overnight. Cells were harvested by centrifugation, resuspended in lysis buffer (20 mM sodium phosphate pH 8.0, 250 mM NaCl, 5 mM imidazole, 0.5 mM PMSF), and lysed by sonication. The lysate was centrifuged (15,000 rpm, 45 min), and the supernatant was filtered and applied to TALON HisPur cobalt resin (Thermo Fisher Scientific). USP7 constructs were eluted with a buffer containing 20 mM sodium phosphate pH 8.0, 250 mM NaCl, and 250 mM imidazole. Thrombin was added to remove the 6-His tag, and the samples were buffer-exchanged overnight at 4 °C into 20 mM sodium phosphate pH 7.4, and 250 mM NaCl to remove imidazole and allow for cleavage. Proteins were then purified by size-exclusion chromatography using a HiLoad Superdex 200 or 75 columns (Cytiva) in a buffer (20 mM Tris pH 7.5, 100 mM NaCl, and 10 mM DTT). Unlabeled ubiquitin used in NMR experiments was expressed in *E. coli* BL21(DE3) cells grown in Luria Broth (LB) media and purified using the same protocol.

USP7 full length and USP7Δtail were expressed transiently in Expi293 (Thermo Fischer Scientific, A14635) following the manufacturer’s manual. Cells were harvested 60-65 hours post-transfection and lysed by sonication in lysis buffer (50 mM HEPES pH 7.4, 200 mM NaCl, 5% glycerol) supplemented with protease inhibitors and benzonase. The lysate was cleared by ultracentrifugation (39,000 rpm, 45 min) and then incubated with FLAG-antibody-coated beads (Genscript, L00432). Bound proteins were eluted with 0.2 mg/mL 1xFLAG (DYKDDDDK) peptide and elution fractions were concentrated. USP7 FL protein was further purified by ion exchange chromatography (IEX) with a Poros 50HQ (Thermo Fisher Scientific, 1255911) column, eluting with a linear NaCl gradient from 200 - 750 mM. Elution fractions were concentrated and polished by size exclusion chromatography (SEC) in SEC buffer (30 mM HEPES pH 7.4, 150 mM NaCl) using a Superdex200Inc 10/300 GL column (Cytiva). IEX was skipped for the purification of USP7Δtail as elutions from FLAG affinity were concentrated and directly injected for SEC.

### Ubiquitin-rhodamine activity assay

DUB activity was tested with a fluorogenic, ubiquitin-rhodamine 110 (UbRho) assay in low-volume, flat-bottom 384-well plates (Corning #3820). Enzyme (USP7cat: 20 nM, USP7Δtail: 20 nM, USP7FL: 10 nM, USP6: 1 nM, USP8: 10 nM, USP28: 10 nM, USP47: 50 nM, UCHL1: 100 nM) was incubated with MS-8 or DMSO for 1 hour at room temperature in assay buffer (50 mM Tris pH 8.0, 50 mM NaCl, 5 mM TCEP, 0.002% Tween20). UbRho (Bio-Techne catalog # U-555-050) was added to a final concentration of 500 nM to initiate the reaction. The enzyme initial velocity was collected over a 30 min period with half minute intervals using a CLARIOstar Microplate Reader (BMG LABTECH) at excitation: 487 nm and emission: 535 nm. The linear regression of the initial velocity (V_0_, RFU/min) was extracted and plotted against MS-8 concentrations using GraphPad Prism (non-linear regression: log(agonist) vs. dose-response (three parameter)). For DUB selectivity assay: initial velocities were normalized to that respective enzyme’s baseline activity with DMSO, and data plotted as fold-change in initial velocity. MS-8 was tested against each enzyme in n = 4, with error bars as SEM.

### Michaelis-Menten Kinetic parameters

Ubiquitin-rhodamine 110 (UbRho) was serially diluted in assay buffer (50 mM Tris pH 8.0, 50 mM NaCl, 5 mM TCEP, 0.002% Tween20) in low-volume, flat-bottom 384-well plates (Corning #3820). USP7 constructs were prepared at 2X final assay concentration in assay buffer and distributed to a 96-well PCR microplate (Axygen #PCR-96-FS-C). MS-8 was serially diluted in the USP7 solution, followed by a 1 hour incubation at room temperature. 10 µL of USP7-MS83 solution was added to 10 µL of UbRho solution in the 384-well plates to initiate the reaction. Fluorescence was measured over 1 hour or 3 hours at half minute intervals using a CLARIOstar Microplate Reader (BMG LABTECH) at excitation: 487 nm and emission: 535 nm. The linear initial velocities were plotted against UbRho concentration for each MS-8 concentration. The RFU standard curve was determined by averaging RFU at saturation (time 2.5 – 3 hour) and plotting against UbRho concentration. The slope of the standard curve (RFU/µM) was used to convert initial reaction rates to µM/min. Kinetic parameters (*k*_cat_ and *K*_M_) for each MS-8 concentration were calculated using GraphPad Prism (Enzyme Kinetics: Velocity as a function of substrate: kcat).

### Ubiquitin-probe labeling assay

*In vitro* USP7 covalent labeling by ubiquitin probe was assessed with compound or DMSO. MS-8 or MS-180 were serially diluted in DMSO, and 2 µL of compound dilution or DMSO were added to the enzyme in PBS buffer (USP7cat: 4 µM or USP7FL: 0.25 µM) in PBS buffer. Ubiquitin-propargylamine (UbiQ, Cat. UbiQ-076) was added to a final concentration of 10X enzyme concentration. Reactions were incubated at room temperature for 2 hours and quenched with 10% DTT in Nu-PAGE LDS Sample Buffer. The USP7 catalytic domain reaction was run on an SDS-PAGE 4-12% gel with Bolt^TM^ MES buffer (Thermo Fisher Scientific) at 175V for 35 mins, and the USP7 full length reaction was run on a NuPAGE^TM^ 3-8% gel (Invitrogen) with Tris-Acetate buffer at 120V for 60 mins, then gels were stained with InstantBlue Coomassie Protein Stain (Abcam # ab119211). The gels were imaged on Amersham Imager 600.

USP7Δtail labeling was assessed using lysates from HCT116 USP7(-/-) cells transfected with USP7Δtail. 2×10^6^ cells were pelleted, then incubated on ice for 30 min with 100 µL of lysis buffer (20 mM Tris pH 8.0, 150 mM NaCl, 10% glycerol, 1% NP-40, 1 mM TCEP) supplemented with protease inhibitors. Lysed cells were centrifuged (15,000 rpm, 4 °C, 10 mins), and the supernatant transferred to new 1.5 mL eppendorf. The lysate protein concentration was determined using a BCA standard titration, then the lysate was diluted to 0.5 mg/ml using lysis buffer. MS-8 dilutions or DMSO were pre-incubated with the lysate at room temperature for 1 hour. A 1:1 mixture of ubiquitin-propargylamine (UbiQ, Cat. UbiQ-076) and ubiquitin-vinylmethylester (Cat. UbiQ-054) was added to a final concentration of 1 µM. Reactions were incubated at room temperature for 2 hours and quenched with 10% DTT in Nu-PAGE LDS Sample Buffer. The NuPAGE^TM^ 3-8% gel (Invitrogen) was run with Tris-Acetate buffer at 120V for 60 mins. Gels were transferred to a PVDF membrane and blocked using 5% non-fat dry milk in 0.1% TBST for 1 hour at room temperature. The membrane was incubated with primary antibodies (anti-HAUSP 1:1,000, anti-β-Actin 1:2,000) overnight at 4 °C, then subsequently 3x washed with 0.1 % TBST. Blots were incubated with secondary antibody (Goat anti-rabbit 1:10,000) for 1 hour at room temperature. Imaging was done in a LI-COR Odyssey CLx 9140 Imaging System, and image processing was performed using Image Studio Lite Ver 5.2.

### Di-ubiquitin cleavage activity assay

USP7 activity was assessed in a di-ubiquitin cleavage assay using K48-linked (LifeSensors), K63-linked (LifeSensors), and linear di-ubiquitin (Ubiquigent). MS-8 dilutions or DMSO were added to 0.2 µM of USP7cat in assay buffer (50 mM Tris pH 8.0, 50 mM NaCl, 5 mM TCEP, 0.002% Tween20). 2 µM of di-ubiquitin was added, and the reaction incubated for 2 hours at 37 °C. Reactions were quenched with 10% BME in LDS Sample Buffer. The SDS-PAGE 4-12% gels were run with Bolt^TM^ MES buffer (Thermo Fisher Scientific) at 175V for 35 mins, then stained with InstantBlue Coomassie Protein Stain (Abcam # ab119211). The gel was destained with milliQ water and imaged on Amersham Imager 600.

### Computational docking and molecular dynamic simulations

We prepared USP7 structures utilizing Protein Preparation Wizard (Schrödinger, Inc.) and prepared the ligand with LigPrep (Schrödinger, Inc.). All the protein-ligand docking has been carried out on the Glide package in Maestro 2022-2.^46^ The inner and outer box sizes are listed in **Supplemental Table 1A**. The conformational sampling with energy window of 2.5 kcal/mol, receptor and ligand van der Waals scaling of 0.5, Cv_cutoff = 100 and Ligand_ccut of 0.15 have been applied. Each binding site defined as the center of the reference ligand on X-ray cocrystal structures. In the docking workflow, molecular docking in *Glide* was first performed, followed by sidechain and loop refinement within 5 Å of MS-8 in *Prime*. Finally, MS-8 was redocked to the refined protein conformations and the Glide XP score was calculated to select the top poses.

All the models were built in Maestro (Schrödinger, Inc.) and prepared for simulation in System Builder. The multistage equilibrium strategy in the Desmond package was used. The simulation box was built with the SPC water model and the OPL3e force field (Roos et al. 2019). All the NPT simulations (300 K, 1 atm) were carried out in the Desmond program (Schrödinger, Inc.) on graphics processing units (GPUs), with a recording interval of 9.6 ps and the van der Waals and short-range electrostatics cut off at 9 Å. Each simulation was performed in triplicate (100 ns each) **(Supplementary Table 1B**). All simulations were analyzed using the Simulation Event Analysis tool implemented in *Maestro.* Molecular visualization is carried out using PyMOL (Version 2.5.4 Schrödinger, LLC).

### NMR Chemical Shift Perturbation

All NMR spectra were collected on an 800MHz (^1^H) Aligent VNMRS spectrometer equipped with a cryoprobe at 30°C. Data processing was performed with NMRPipe,^45^ and analysis was done using Sparky.^44^ NMR buffer contained 20 mM Tris pH 7.5, 100 mM NaCl, 2 mM DTT, and 10% D_2_O (v/v).

To test activator binding to WT USP7 a 20 mM stock of MS-8 in 100% DMSO was gradually added to 0.25 mM ^15^N WT USP7 catalytic domain to a molar ratio of 1:4 (protein:activator). ^1^H-^15^N TROSY spectra of USP7 were collected at each of the 6 titration points. DMSO was used as a negative control. The spectra were analyzed in Sparky and chemical shift perturbations (CSPs), Δω, caused by the addition of MS-8 were calculated for every peak in the spectrum using the following equation:

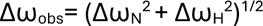

where Δω_N_ and Δω_H_ are the frequency differences between free and ligand-bound samples measured in Hz for ^15^N and ^1^H, respectively. The frequency perturbations were converted to chemical shift perturbations by dividing Δω_obs_ by ^1^H frequency of the spectrometer. The Δω values were mapped onto the crystal structure of USP7 (PDB ID: 4M5W) using PyMOL.

To test MS-8 binding to C223A alone or its complex with ubiquitin (C223A/ubiquitin), a 10 mM stock of MS-8 in 100% DMSO was gradually added to 0.3 mM ^15^N C223A (or C223A/ubiquitin) up to a 4-fold molar excess of MS-8. C223A/Ubiquitin complex was obtained by mixing ^15^N C223A USP7 catalytic domain with the 4-fold excess of unlabeled ubiquitin and following the binding saturation using ^1^H-^15^N TROSY NMR spectra. The binding of MS-8 to C223A and C223A/ubiquitin was monitored at each of the titration points using ^1^H-^15^N TROSY spectra. DMSO was used as a negative control. The Δω values were determined for all peaks in the spectrum at each titration point. Perturbations of peaks corresponding to residues in the CBP pocket (K281, S282 and G284) as a function of MS-8 concentration were used to illustrate the binding curves using one site-specific binding model in GraphPad Prism (version 10.4.1).

### Fluorescence polarization (FP) binding assay

USP7 constructs were serial diluted in FP assay buffer (20 mM HEPES pH 7.5, 100 mM NaCl, 1 mM DTT, 1 mM EDTA, 0.05% Tween20) in a low-volume, flat-bottom 384-well plates (Corning #3820). DMSO or MS-8 were added to the enzyme. N-terminally tetramethylrhodamine (TAMRA)- labeled ubiquitin (LifeSensors, SI270T) was added to all the assay wells. The mixture was incubated for 2 hours at room temperature. Fluorescence polarization was measured on a PHERAstar *FS* (BMB LABTECH) at excitation 540 nm and emission 590 nm, calibrated to a gain of 35 mA. Changes in anisotropy were calculated on the MARS Analysis Software (BMG LABTECH) and plotted versus concentration of enzyme using GraphPad Prism.

Competition FP was measured using N-terminally TAMRA-USP7 C-terminal domains (CTD). TAMRA-probe was generated by expression with pGEX6P-USP7(883-end), purification as described for USP7 catalytic domain, followed by 5-TAMRA labeling and purification using the AnaTag^TM^ 5-TAMRA Protein Labeling Kit (AnaSpec, Inc.). USP7 catalytic domain (250 µM in FP assay buffer) was pre-incubated with MS-8 dilutions or DMSO for 1 hour at room temperature, followed by addition of TAMRA-USP7_CTD_ (1 µM in FP assay buffer). The mixture was incubated for 2 hours at room temperature. Fluorescence polarization was measured on a PHERAstar *FS* (BMB LABTECH) at excitation 540 nm and emission 590 nm, calibrated to a gain of 35 mA. Changes in anisotropy were calculated on the MARS Analysis Software (BMG LABTECH) and plotted versus concentration of MS-8 using GraphPad Prism.

### General Cell Culture

HCT 116 cells (ATCC® CCL-247™) were grown in McCoy’s 5A Medium (ATCC® 30-2007™) supplemented with 10% fetal bovine serum (FBS) in a humidified 37°C incubator at 5% CO_2_. HCT 116 USP7(-/-) line was purchased from Horizon Discovery (HD R02-028) and cultured following the same conditions as the parental line.

Resuspension of adherent cells was conducted by vacuum aspirating the media and replacing it with warmed supplemented media using an electronic pipette gun. Cell suspension was subsequently collected and pelleted by centrifugation (300 x g, 5 min). Cell concentration was determined by using a Countess® II FL Automated Cell Counter, and cells were plated accordingly.

### Luminescent Cell Viability Assay

The CellTiter Glo® (Promega) viability assay was performed by plating HCT 116 USP7(+/+) and USP7(-/-) cells at a density of 1 x 10^4^ cells/well in a clear-bottom, white 96-well plate (Corning #3903) (100 μL total volume). The plate was placed in the cell culture incubator overnight to allow full attachment to the wells. Next day, media from all wells was removed using a multichannel pipette, and stock solutions of 100 μL of either MS-8 or MS-180 in DMSO were prepared and subsequently added in triplicate to the wells. (Final concentration of DMSO of 1%). After allowing incubation for 24 hours in the culture incubator, the plate was taken out and allowed to reach room temperature for 30 min. CellTiter-Glo® mix (prepared per manufacturer’s protocol) was thawed and equilibrated to room temperature and subsequently added (50 μL) to each well. The plate was covered with aluminum foil and shaken gently for 10 min in an orbital shaker before reading luminescent signal on a CLARIOstar Microplate Reader (BMG LABTECH).

### Immunoblotting assay

HCT 116 USP7(+/+) and USP7(-/-) cells were plated in a 6-well plate at a density of 800.000 cells/well (2 mL of total volume). The plate was placed in the culture incubator overnight to allow complete attachment to the bottom of the wells. The next day, media was vacuum aspirated, and cells were treated with MS-8, MS-180, XL177A or DMSO (1%) for 24 hours. Media was subsequently vacuum aspirated, washed with PBS and lysed using M-PER lysis buffer (Thermo Fisher Scientific) + 100x HALT protease inhibitor cocktail (Thermo Fisher Scientific). Lysates were clarified by centrifugation (14.000 rpm, 4 °C, 10 min). Total protein concentration was measured using a NanoDrop^TM^ 2000 spectrophotometer. Lysate samples were prepared using 4x SDS Loading buffer + 20x DTT and separated by gel electrophoresis by loading the samples in 4 – 12 % Bolt^TM^ Bis-Tris Plus Mini Protein Gels in a Mini Gel Tank chamber at 120V for 60 min. Gels were transferred to a PVDF membrane and blocked using 5% non-fat dry milk in 0.1% TBST for 1 hour at room temperature. Membranes were probed with the corresponding primary antibodies overnight at 4 °C (1:1,000 for all except anti-TRIM27 1:500). Blots were subsequently 3x washed with 0.1 % TBST and incubated with secondary antibody (1:10,000) for 1 hour at room temperature. Imaging was done in a LI-COR Odyssey CLx 9140 Imaging System, and image processing was performed using Image Studio Lite Ver 5.2.

Analysis of transfected USP7(-/-) cells with USP7Δtail and WT constructs was conducted by mixing 2 ug of vector and Lipofectamine^TM^ 3000 (Invitrogen) in Opti-MEM^TM^ (Gibco) medium not supported with FBS and allowing the cells to grow for 48 hours before dosing with small molecules or DMSO.

### MS-8 synthesis and characterization

All commercially available starting materials were purchased from Sigma Aldrich, Fisher Scientific, Oakwood Chemical and Combi Blocks, and anhydrous solvents were purchased from Fisher Scientific. All reagents and solvents were used as received without further purification. If necessary, air or moisture sensitive reactions were carried out under an inert nitrogen atmosphere. Removal of solvents was accomplished on a Büchi R-300 rotary evaporator connected to a Welch 1400B-01 vacuum line and Labconco FreeZone 6 plus system. Compound purification was achieved using normal phase column chromatography (Teledyne CombiFlash chromatography system) and/or reversed phase chromatography (Shimadzu system with SunFire® Prep C18 OBDTM 5μM column). Purity was assessed by UPLC (Waters Acquity system) and analytical thin layer chromatography (TLC, EMD Millipore TLC Silica Gel60 F254). TLC visualization was accomplished by irradiation under UV light (254nm). All ^1^H-NMR spectra were recorded at 298K on a Bruker ARX 500 (500MHz) spectrometer, and all ^13^C-NMR spectra were recorded on a Bruker ARX 500 (126 MHz) spectrometer. Samples were dissolved in CDCl_3_, DMSO-d_6_, or CD_3_OD, and spectra were referenced to the residual solvent peak (chloroform-d: 7.26 ppm for ^1^H-NMR and 77.16 ppm for ^13^C-NMR; DMSO-d_6_: 2.50 ppm for ^1^H-NMR and 39.25 ppm for ^13^C-NMR, CD_3_OD: 3.31 ppm (-Me) for ^1^H NMR and 49.00 ppm for ^13^C-NMR or tetramethylsilane (TMS) as the internal standard. NMR peaks are reported below with chemical shift, multiplicity (s=singlet, d=doublet, t=triplet, q=quartet, m=multiplet, br=broad peak), coupling constants (Hz), and number of protons. Mass spectrometry (LCMS) data were obtained on Waters Acquity UPLC system in positive ESI mode.

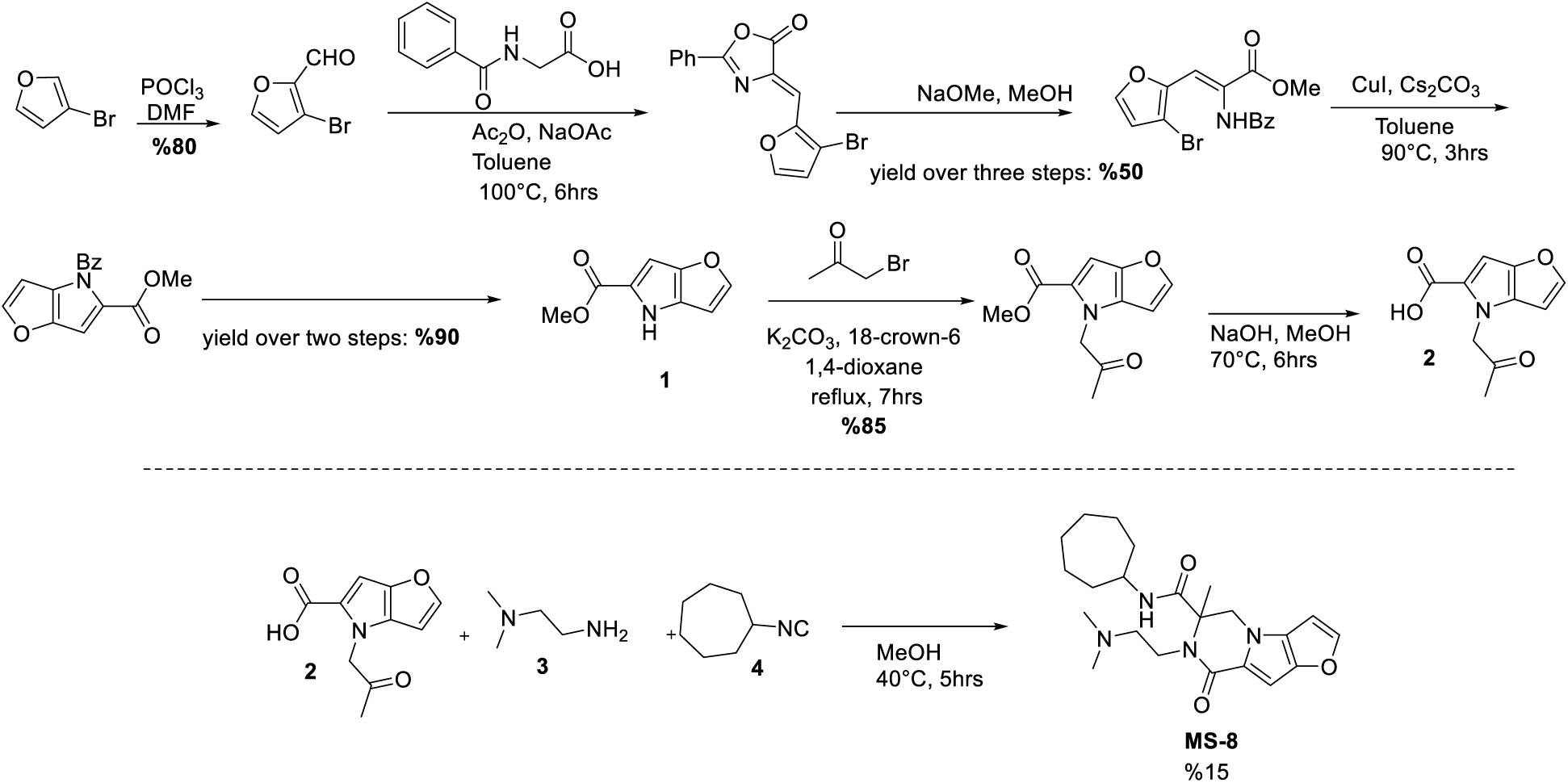

Scheme 1. Synthesis of MS-8.

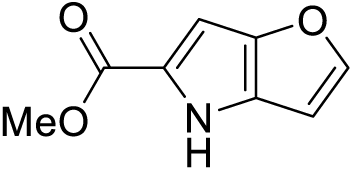

Compound **1** was generated following the previously reported procedure.^47^ ^1^H NMR (500 MHz, DMSO) δ 11.69 (s, 1H), 7.79 (d, *J* = 1.9 Hz, 1H), 6.76 (s, 1H), 6.61 (d, *J* = 2.5 Hz, 1H), 3.79 (s, 3H). ^13^C NMR (125 MHz, DMSO) δ 162.1, 149.8, 147.5, 129.8, 123.7, 99.9, 96.5, 51.7. The NMR data is consistent with literature values.^47^

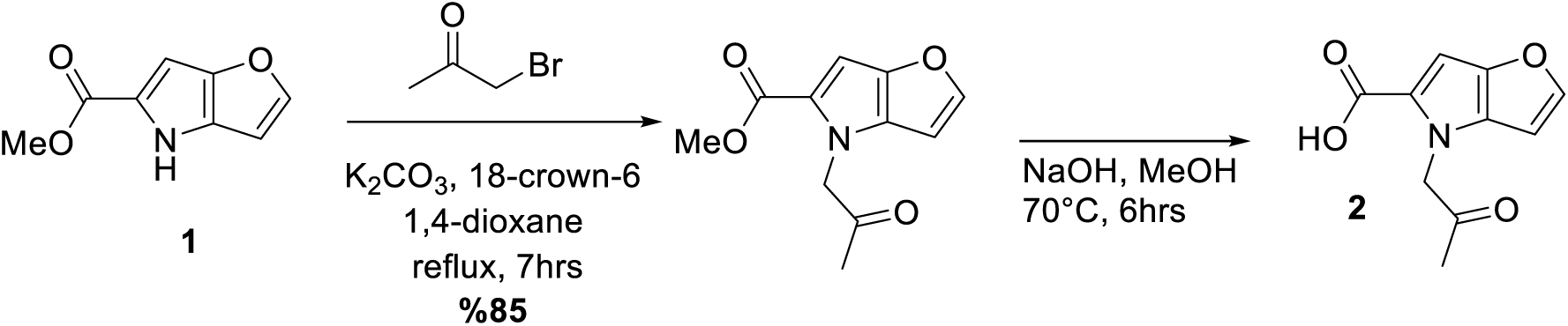

Compound **2** was prepared following previous literature preparation.^48^ **1** (1 eq), α-bromo ketone (1 eq), K_2_CO_3_ (1.2 eq), and 18-crown-6 phase-transfer catalyst (0.1 eq) in 1,4-dioxane (0.2M) was stirred at reflux for 7 h. The reaction mixture was cooled, diluted with DCM and quenched with water. The aqueous layer was then extracted twice with DCM (20 mL). The combined DCM layers were dried with Na_2_SO_4_ and concentrated in vacuo. The resulting oil was used without further purification and used directly in the next reaction. The resulting oil was dissolved in 2:1 methanol: water and a solution of 1M NaOH was added (10 eq.) The reaction was stirred for 6h at 70°C. At completion of the reaction, methanol was removed en vacuo and the pH was adjusted to 2 using 1M HCl. The resulting precipitate **2** was collected by filtration. ^1^H NMR (500 MHz, DMSO) δ 7.79 (d, *J* = 2.3 Hz, 1H), 6.81 (s, 1H), 6.72 (s, 1H), 5.24 (s, 2H), 2.13 (s, 3H). ^13^C NMR (125 MHz, DMSO) δ 203.2, 163.2, 149.4, 145.2, 133.6, 124.2, 99.6, 98.5, 56.8, 27.2.

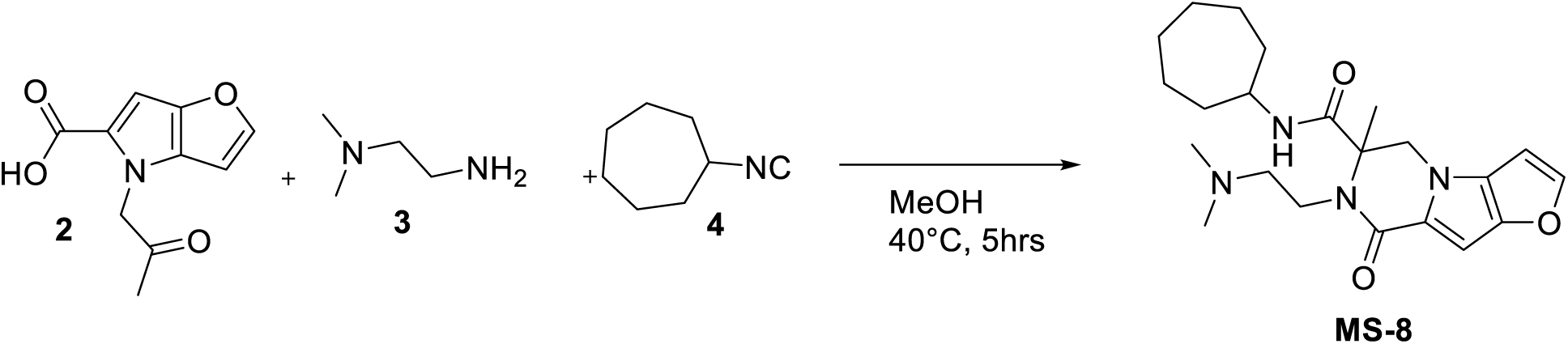

Compound **2** (1 eq) was dissolved in MeOH, compounds **3** (1 eq) and **4** (1 eq) were added and the reaction mixture was heated to 40°C for 5 hours. The solvent was removed, and the crude mixture was purified by HPLC to yield pure **MS-8**.

^1^H NMR (500 MHz, DMSO) δ 8.22 (d, *J* = 7.8 Hz, 1H), 7.73 (d, *J* = 2.2 Hz, 1H), 6.68 (dd, *J* = 2.3, 0.9 Hz, 1H), 6.58 (d, *J* = 0.8 Hz, 1H), 4.67 (d, *J* = 12.4 Hz, 1H), 4.02 (d, *J* = 12.4 Hz, 1H), 3.72 (m, 1H), 3.65 – 3.43 (m, 2H), 2.48 – 2.38 (m, 2H), 2.23 (s, 6H), 1.60 (s, 3H), 1.52 – 1.23 (m, 12H).

^13^C NMR (125 MHz, DMSO) δ 170.7, 160.7, 148.7, 147.1, 128.4, 126.5, 99.3, 93.6, 64.8, 59.1, 51.9, 51.2, 46.0, 34.3, 28.0, 27.8, 24.4, 24.2, 20.6. (2 Carbon peaks covered by DMSO solvent peak) MS-12225-83 (MS-8): LC-MS (ESI) [M+H]^+^ calcd for C22H32N4O3+ m/z 401.3, found m/z 401.31

### MS-180 synthesis and characterization

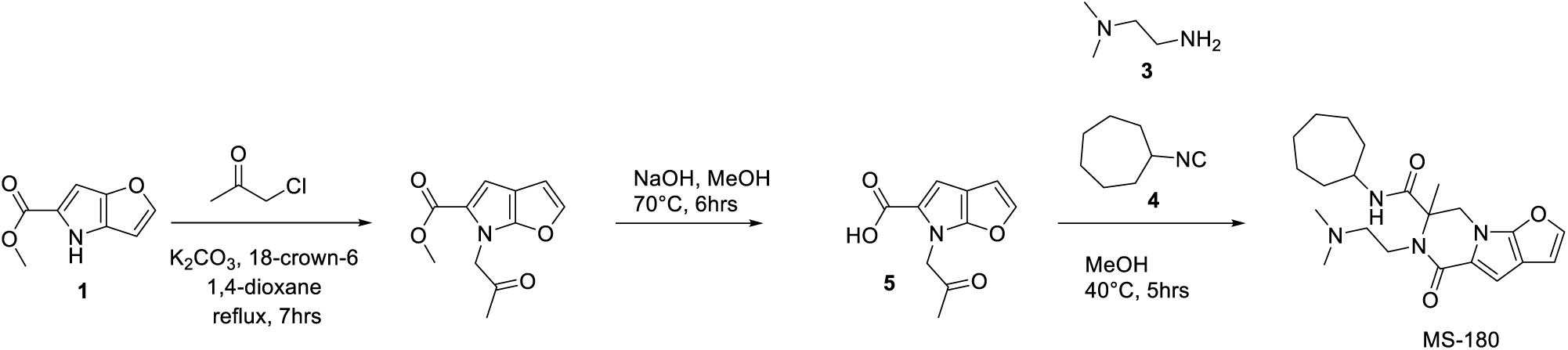

MS-180 was synthesized according to Scheme 2. MS-180 was generated using the same preparations as described for MS-8 above, following previous literature preparation.^47,48^

^1^H NMR (500 MHz, DMSO) δ 7.62 (d, *J* = 2.4 Hz, 1H), 6.67 (s, 1H), 6.64 (d, *J* = 2.0 Hz, 1H), 4.91 (d, *J* = 13.0 Hz, 1H), 4.17 (d, *J* = 13.0 Hz, 1H), 3.90 (dt, *J* = 14.3, 6.7 Hz, 1H), 3.72 (m, 1H), 3.58 (m, 1H), 3.23 (q, *J* = 6.0 Hz, 2H), 2.89 (d, *J* = 4.9 Hz, 6H), 1.69 (s, 3H), 1.54 (ddd, *J* = 30.5, 12.4, 6.2 Hz, 3H), 1.42 (dt, *J* = 13.6, 8.7 Hz, 3H), 1.32 (dd, *J* = 13.3, 6.4 Hz, 4H), 1.17 (dt, *J* = 15.1, 9.0 Hz, 2H).

^13^C NMR (126 MHz, DMSO) δ 170.1, 162.2, 149.6, 145.1, 122.5, 109.7, 106.4, 102.6, 65.3, 57.1, 51.2, 50.1, 43.5, 37.8, 34.3, 34.3, 28.2, 28.1, 24.3, 23.9, 20.4.

MS-13377-180 (MS-180): LC-MS (ESI) [M+H]^+^ calcd for C22H33N4O3+ m/z 401.25, found m/z 401.3

## ABBREVIATIONS

USP7: Ubiquitin-specific protease 7
HAFOUS: Hao-Fountain Syndrome
DUB: Deubiquitylating enzyme, Deubiquitinase
Ub: Ubiquitin
UBL: Ubiquitin-like domain
TRAF: Tumor necrosis factor receptor-associated factors
CTD: C-terminal domain
CPB: C-terminal peptide binding
MMGBSA: Molecular Mechanics with Generalized Born and Surface Area
RMSD: Root mean square deviation
MD: Molecular dynamic
FP: Fluorescence polarization
TAMRA: tetramethylrhodamine
Ub-PA: ubiquitin-propargylamine
Ub-VME: ubiquitin-vinylmethyl ester
UbRho: Ubiquitin-rhodamine 110

---

**Supplemental Figure 1**

(**A**) Initial screening data for MS-8 against eight recombinant DUBs in ubiquitin-rhodamine assay. Full screening information in Varca et al. Ubiquitin-rhodamine activity assay raw fluorescence (RFU) over time (min) with MS-8 in dose response against (**B**) USP7 catalytic domain and (**C**) USP7 full length. Fold change with dose-response MS-8 calculated from initial velocities with activator divided by initial velocity with DMSO. Fracture in signal at the 60-minute mark for all concentrations and both enzymes due to instrument error. Comparison between catalytic domain and full length as Initial Velocity/[Enzyme] in Figure 1B. (**D**) Comparison between separated enantiomers of MS-8 against USP7 catalytic domain, showing a preference for MS-8-P1. n = 2. (**E**) Ubiquitin-propargylamine (Ub-PA) *in vitro* labeling gel, with USP7 catalytic domain with MS-8 enantiomers. USP7 catalytic domain was pre-incubated with MS-8 for 1 hour, followed by a 2 hour reaction with Ub-PA at room temperature.

**Supplemental Figure 2**

(**A**) Chemical structure of MS-180, negative control of MS-8. (**B**) Comparison between negative control (MS-180) and activator (MS-8) in fluorogenic ubiquitin-rhodamine assay, using catalytic domain USP7. (**C**) Ubiquitin-propargylamine labeling of USP7 catalytic domain with MS-180 dosing. Reaction incubated for 2 hours. (**D**) Di-ubiquitin cleavage with USP7 catalytic domain and MS-180 dosing. K48- and K63-linked di-ubiquitin were each assessed. (**E**) MS-180 dose-response ubiquitin-rhodamine assay with USP7 catalytic domain mutants, with mutations in C-terminal peptide binding pocket. Fold change with dose-response MS-180 calculated from initial velocities with compound divided by initial velocity with DMSO. n = 4. (**F**) MS-180 with USP7 catalytic domain, USP7Δtail, and USP7 full length. n = 4.

**Supplemental Figure 3**

Michaelis-Menten kinetic assay with MS-8 and (**A**) USP7 catalytic domain, (**B**) USP7Δtail, (**C**) USP7 full length. USP7 catalytic domain and USP7Δtail were tested at 20 nM, and USP7 full length at 10 nM.

**Supplemental Figure 4**

(**A**) Overlay of ^15^N TROSY NMR spectra of the USP7 catalytic domain in complex with MS-8 (black) and in the presence of 4% DMSO (red). Insets highlight two regions (outlined by black boxes) corresponding to the amide peaks of residues K281 and W285, located within the C-terminal peptide binding (CPB) pocket of the USP7 catalytic domain. (**B**) Chemical shift perturbations (Δω) from panel A mapped on the surface of USP7 catalytic domain (PDB ID: 4M5W) and colored from white (smallest Δω) to pink (largest Δω). MS-8 (green) posed from molecular dynamics simulation is overlaid with the Δω-mapped structure. Inset shows a zoomed-in region of the C-terminal peptide binding pocket of USP7. (**C**) Overlay of ^15^N TROSY NMR spectra of the USP7^C223A^ catalytic domain from Figure 4B-C. Red – free USP7 in the presence of 5% DMSO; blue – USP7 in complex with ubiquitin (1:4 molar ratio) in the presence of 5% DMSO; black – USP7 in complex with ubiquitin and 4-molar excess of MS-8.

**Supplemental Figure 5**

Cell viability with (**A**) activator MS-8 and (**B**) negative control MS-180 in both HCT116 parental USP7(+/+) and HCT116 USP7 (-/-) cells. Cells were incubated with small molecules for 24 hours. n = 3. (**C**) Full western blots of Figure 5A, with box around section that appears in the main paper. TRIM27 and αTubulin were imaged from the same section of blot, but using the 800 nm and 680 nm channels, respectively. (**D-E**). Full western blots of Figure 5B, with box around section that appears in the main paper.

**Supplemental Figure 6**

Enzyme titrations of USP7 catalytic domain mutants assessed using ubiquitin-rhodamine at 500 µM. n = 2.

**Supplemental Figure 7**

Di-ubiquitin cleavage full gels, with box around section that appears in main paper figures.

**Supplemental Figure 8**

Covalent ubiquitin labeling full gels. (**A-E**) Gels from in vitro ubiquitin-propargylamine (Ub-PA) labeling were Coomassie stained and imaged. (**F**) Gel from lysate labeling with 1:1 mixture of ubiquitin-propargylamine (Ub-PA):ubiquitin-vinylmethylester (Ub-VME). Western blot with 1:1000 1° anti-USP7 Rb, 1:2000 1° anti-β-Actin Rb, and 1:10,000 2° Goat anti-Rb.

**Supplemental Table 1**

(**A**) Docking score (Glide XP precision) of the top pose for each pocket with MS-8. GNE-6776 was docked into distal pocket as control. (**B**) Summary of all-atom molecular dynamic simulations.

**Supplemental Table 2**

(**A**) Forward and reverse primer sequences for USP7 catalytic domain mutants. (B) Forward and reverse primer sequences for USP7Δtail mutant. (**C**) Dissociation constant (KD) from Figure 4A. Values derived from GraphPad Prism: log(agonist) vs response (three parameter). USP7 catalytic domain + 100 µM MS-8, n = 1, all others n = 2.

## REFERENCES

1. Callis, J. (2014). The Ubiquitination Machinery of the Ubiquitin System. Arabidopsis Book 12, e0174. 10.1199/tab.0174.

2. Bonacci, T., and Emanuele, M.J. (2020). Dissenting degradation: Deubiquitinases in cell cycle and cancer. Semin Cancer Biol 67, 145–158. 10.1016/j.semcancer.2020.03.008.

3. Snyder, N.A., and Silva, G.M. (2021). Deubiquitinating enzymes (DUBs): Regulation, homeostasis, and oxidative stress response. Journal of Biological Chemistry 297. 10.1016/j.jbc.2021.101077.

4. Pozhidaeva, A., and Bezsonova, I. (2019). USP7: Structure, substrate specificity, and inhibition. DNA Repair (Amst) 76, 30–39. 10.1016/j.dnarep.2019.02.005.

5. Lamberto, I., Liu, X., Seo, H.S., Schauer, N.J., Iacob, R.E., Hu, W., Das, D., Mikhailova, T., Weisberg, E.L., Engen, J.R., et al. (2017). Structure-Guided Development of a Potent and Selective Non-covalent Active-Site Inhibitor of USP7. Cell Chem Biol 24, 1490–1500.e11. 10.1016/j.chembiol.2017.09.003.

6. Schauer, N.J., Liu, X., Magin, R.S., Doherty, L.M., Chan, W.C., Ficarro, S.B., Hu, W., Roberts, R.M., Iacob, R.E., Stolte, B., et al. (2020). Selective USP7 inhibition elicits cancer cell killing through a p53-dependent mechanism. Sci Rep 10. 10.1038/s41598-020-62076-x.

7. Al-Eidan, A., Wang, Y., Skipp, P., and Ewing, R.M. (2022). The USP7 protein interaction network and its roles in tumorigenesis. Genes Dis 9, 41–50. 10.1016/j.gendis.2020.10.004.

8. Kon, N., Zhong, J., Kobayashi, Y., Li, M., Szabolcs, M., Ludwig, T., Canoll, P.D., and Gu, W. (2011). Roles of HAUSP-mediated p53 regulation in central nervous system development. Cell Death Differ 18, 1366–1375. 10.1038/cdd.2011.12.

9. Hao, Y.H., Fountain, M.D., Fon Tacer, K., Xia, F., Bi, W., Kang, S.H.L., Patel, A., Rosenfeld, J.A., Le Caignec, C., Isidor, B., et al. (2015). USP7 Acts as a Molecular Rheostat to Promote WASH-Dependent Endosomal Protein Recycling and Is Mutated in a Human Neurodevelopmental Disorder. Mol Cell 59, 956–969. 10.1016/j.molcel.2015.07.033.

10. Sijm, A., Atlasi, Y., van der Knaap, J.A., Wolf van der Meer, J., Chalkley, G.E., Bezstarosti, K., W Dekkers, D.H., S Doff, W.A., Ozgur, Z., J van IJcken, W.F., et al. (2022). USP7 regulates the ncPRC1 Polycomb axis to stimulate genomic H2AK119ub1 deposition uncoupled from H3K27me3.

11. Lecona, E., Narendra, V., and Reinberg, D. (2015). USP7 Cooperates with SCML2 To Regulate the Activity of PRC1. Mol Cell Biol 35, 1157–1168. 10.1128/mcb.01197-14.

12. Fountain, M.D., Oleson, D.S., Rech, M.E., Segebrecht, L., Hunter, J. V, McCarthy, J.M., Lupo, P.J., Holtgrewe, M., Moran, R., Rosenfeld, J.A., et al. Pathogenic variants in USP7 cause a neurodevelopmental disorder with speech delays, altered behavior, and neurologic anomalies. GENETICS in MEDICINE. 10.1038/s41436.

13. Faesen, A.C., Dirac, A.M.G., Shanmugham, A., Ovaa, H., Perrakis, A., and Sixma, T.K. (2011). Mechanism of USP7/HAUSP activation by its C-Terminal ubiquitin-like domain and allosteric regulation by GMP-synthetase. Mol Cell 44, 147–159. 10.1016/j.molcel.2011.06.034.

14. Kim, R.Q., Geurink, P.P., Mulder, M.P.C., Fish, A., Ekkebus, R., El Oualid, F., van Dijk, W.J., van Dalen, D., Ovaa, H., van Ingen, H., et al. (2019). Kinetic analysis of multistep USP7 mechanism shows critical role for target protein in activity. Nat Commun 10. 10.1038/s41467-018-08231-5.

15. Özen, A., Rougé, L., Bashore, C., Hearn, B.R., Skelton, N.J., and Dueber, E.C. (2018). Selectively Modulating Conformational States of USP7 Catalytic Domain for Activation. Structure 26, 72–84.e7. 10.1016/j.str.2017.11.010.

16. Rougé, L., Bainbridge, T.W., Kwok, M., Tong, R., Di Lello, P., Wertz, I.E., Maurer, T., Ernst, J.A., and Murray, J. (2016). Molecular Understanding of USP7 Substrate Recognition and C-Terminal Activation. Structure 24, 1335–1345. 10.1016/j.str.2016.05.020.

17. Schauer, N.J., Magin, R.S., Liu, X., Doherty, L.M., and Buhrlage, S.J. (2020). Advances in Discovering Deubiquitinating Enzyme (DUB) Inhibitors. J Med Chem 63, 2731–2750. 10.1021/acs.jmedchem.9b01138.

18. Turnbull, A.P., Ioannidis, S., Krajewski, W.W., Pinto-Fernandez, A., Heride, C., Martin, A.C.L., Tonkin, L.M., Townsend, E.C., Buker, S.M., Lancia, D.R., et al. (2017). Molecular basis of USP7 inhibition by selective small-molecule inhibitors. Nature 550, 481–486. 10.1038/nature24451.

19. Gavory, G., O’Dowd, C.R., Helm, M.D., Flasz, J., Arkoudis, E., Dossang, A., Hughes, C., Cassidy, E., McClelland, K., Odrzywol, E., et al. (2018). Discovery and characterization of highly potent and selective allosteric USP7 inhibitors. Nat Chem Biol 14, 118–125. 10.1038/nchembio.2528.

20. Li, X., Yang, S., Zhang, H., Liu, X., Gao, Y., Chen, Y., Liu, L., Wang, D., Liang, Z., Liu, S., et al. (2022). Discovery of Orally Bioavailable N-Benzylpiperidinol Derivatives as Potent and Selective USP7 Inhibitors with in Vivo Antitumor Immunity Activity against Colon Cancer. J Med Chem 65, 16622–16639. 10.1021/acs.jmedchem.2c01444.

21. Ohol, Y.M., Sun, M.T., Cutler, G., Leger, P.R., Hu, D.X., Biannic, B., Rana, P., Cho, C., Jacobson, S., Wong, S.T., et al. (2020). Novel, selective inhibitors of USP7 uncover multiple mechanisms of antitumor activity in vitro and in vivo. Mol Cancer Ther 19, 1970–1980. 10.1158/1535-7163.MCT-20-0184.

22. Pei, Y., Fu, J., Shi, Y., Zhang, M., Luo, G., Luo, X., Song, N., Mi, T., Yang, Y., Li, J., et al. (2022). Discovery of a Potent and Selective Degrader for USP7. Angewandte Chemie - International Edition 61. 10.1002/anie.202204395.

23. Murgai, A., Sosič, I., Gobec, M., Lemnitzer, P., Proj, M., Wittenburg, S., Voget, R., Gütschow, M., Krönke, J., and Steinebach, C. (2022). Targeting the deubiquitinase USP7 for degradation with PROTACs. Chemical Communications 58, 8858–8861. 10.1039/d2cc02094g.

24. Liu, J., Hu, X., Luo, K., Xiong, Y., Chen, L., Wang, Z., Inuzuka, H., Qian, C., Yu, X., Xie, L., et al. (2024). USP7-Based Deubiquitinase-Targeting Chimeras Stabilize AMPK. J Am Chem Soc. 10.1021/jacs.4c02373.

25. Huang, Y.T., Cheng, A.C., Tang, H.C., Huang, G.C., Cai, L., Lin, T.H., Wu, K.J., Tseng, P.H., Wang, G.G., and Chen, W.Y. (2021). USP7 facilitates SMAD3 autoregulation to repress cancer progression in p53-deficient lung cancer. Cell Death Dis 12. 10.1038/s41419-021-04176-8.

26. Frey, Y., Franz-Wachtel, M., Macek, B., and Olayioye, M.A. (2022). Proteasomal turnover of the RhoGAP tumor suppressor DLC1 is regulated by HECTD1 and USP7. Sci Rep 12. 10.1038/s41598-022-08844-3.

27. Varca, A.C., Casalena, D., Chan, W.C., Hu, B., Magin, R.S., Roberts, R.M., Liu, X., Zhu, H., Seo, H.S., Dhe-Paganon, S., et al. (2021). Identification and validation of selective deubiquitinase inhibitors. Cell Chem Biol 28, 1758–1771.e13. 10.1016/j.chembiol.2021.05.012.

28. Valles, G.J., Bezsonova, I., Woodgate, R., and Ashton, N.W. (2020). USP7 Is a Master Regulator of Genome Stability. Front Cell Dev Biol 8. 10.3389/fcell.2020.00717.

29. Kategaya, L., Di Lello, P., Rougé, L., Pastor, R., Clark, K.R., Drummond, J., Kleinheinz, T., Lin, E., Upton, J.P., Prakash, S., et al. (2017). USP7 small-molecule inhibitors interfere with ubiquitin binding. Nature 550, 534–538. 10.1038/nature24006.

30. Morrow, M.E., Morgan, M.T., Clerici, M., Growkova, K., Yan, M., Komander, D., Sixma, T.K., Simicek, M., and Wolberger, C. (2018). Active site alanine mutations convert deubiquitinases into high-affinity ubiquitin-binding proteins. EMBO Rep 19. 10.15252/embr.201745680.

31. Dow, L.F., Case, A.M., Paustian, M.P., Pinkerton, B.R., Simeon, P., and Trippier, P.C. (2023). The evolution of small molecule enzyme activators. RSC Med Chem 14, 2206–2230. 10.1039/d3md00399j.

32. Teng, M., Young, D.W., and Tan, Z. (2022). The Pursuit of Enzyme Activation: A Snapshot of the Gold Rush. J Med Chem 65, 14289–14304. 10.1021/acs.jmedchem.2c01291.

33. Xiao, B., Sanders, M.J., Carmena, D., Bright, N.J., Haire, L.F., Underwood, E., Patel, B.R., Heath, R.B., Walker, P.A., Hallen, S., et al. (2013). Structural basis of AMPK regulation by small molecule activators. Nat Commun 4. 10.1038/ncomms4017.

34. Calabrese, M.F., Rajamohan, F., Harris, M.S., Caspers, N.L., Magyar, R., Withka, J.M., Wang, H., Borzilleri, K.A., Sahasrabudhe, P. V., Hoth, L.R., et al. (2014). Structural basis for AMPK activation: Natural and synthetic ligands regulate kinase activity from opposite poles by different molecular mechanisms. Structure 22, 1161–1172. 10.1016/j.str.2014.06.009.

35. Grimsby, J., Sarabu, R., Corbett, W.L., NHaynes, N.-E., Bizzarro, F.T., Coffey, J.W., Guertin, K.R., Hilliard, D.W., Kester, R.F., Mahaney, P.E., et al. (2003). Allosteric activators of glucokinase: potential role in diabetes therapy. Science (1979) 301, 370–373.

36. Tawa, P., Zhang, L., Metwally, E., Hou, Y., McCoy, M.A., Seganish, W.M., Zhang, R., Frank, E., Sheth, P., Hanisak, J., et al. (2022). Mechanistic insights on novel small molecule allosteric activators of cGMP-dependent protein kinase PKG1α. Journal of Biological Chemistry 298. 10.1016/j.jbc.2022.102284.

37. Kok, B.P., Ghimire, S., Kim, W., Chatterjee, S., Johns, T., Kitamura, S., Eberhardt, J., Ogasawara, D., Xu, J., Sukiasyan, A., et al. (2020). Discovery of small-molecule enzyme activators by activity-based protein profiling. Nat Chem Biol 16, 997–1005. 10.1038/s41589-020-0555-4.

38. Gong, G.Q., Bilanges, B., Allsop, B., Masson, G.R., Roberton, V., Askwith, T., Oxenford, S., Madsen, R.R., Conduit, S.E., Bellini, D., et al. (2023). A small-molecule PI3Kα activator for cardioprotection and neuroregeneration. Nature 618, 159–168. 10.1038/s41586-023-05972-2.

39. Shi, L., Xu, Z., Chen, X., Meng, Q., Zhou, H., Xiong, B., and Zhang, N. (2025). Sertraline and Astemizole Enhance the Deubiquitinase Activity of USP7 by Binding to Its Switching Loop Region. J Med Chem. 10.1021/acs.jmedchem.5c00032.

40. Pilarski, R., Carlo, M.I., Cebulla, C., and Abdel-Rahman, M. (2016). BAP1 Tumor Predisposition Syndrome. https://www.ncbi.nlm.nih.gov/books/NBK390611/.

41. Lu, D., Song, J., Sun, Y., Qi, F., Liu, L., Jin, Y., McNutt, M.A., and Yin, Y. (2018). Mutations of deubiquitinase OTUD1 are associated with autoimmune disorders. J Autoimmun 94, 156–165. 10.1016/j.jaut.2018.07.019.

42. Henning, N.J., Boike, L., Spradlin, J.N., Ward, C.C., Liu, G., Zhang, E., Belcher, B.P., Brittain, S.M., Hesse, M.J., Dovala, D., et al. (2022). Deubiquitinase-targeting chimeras for targeted protein stabilization. Nat Chem Biol 18, 412–421. 10.1038/s41589-022-00971-2.

43. Pozhidaeva, A., Valles, G., Wang, F., Wu, J., Sterner, D.E., Nguyen, P., Weinstock, J., Kumar, K.G.S., Kanyo, J., Wright, D., et al. (2017). USP7-Specific Inhibitors Target and Modify the Enzyme’s Active Site via Distinct Chemical Mechanisms. Cell Chem Biol 24, 1501–1512.e5. 10.1016/j.chembiol.2017.09.004.

44. T. D. Goddard, and D. G. Kneller (2000). SPARKY 3. https://www.cgl.ucsf.edu/home/sparky/.

45. Delaglio ∼’, F., Grzesiek, S., Vuister, G.W., Zhu, G., Pfeifer, J., and Bax, A. (1995). NMRPipe: A multidimensional spectral processing system based on UNIX pipes*. J Biomol NMR 6, 293.

46. Friesner, R.A., Murphy, R.B., Repasky, M.P., Frye, L.L., Greenwood, J.R., Halgren, T.A., Sanschagrin, P.C., and Mainz, D.T. (2006). Extra precision glide: Docking and scoring incorporating a model of hydrophobic enclosure for protein-ligand complexes. J Med Chem 49, 6177–6196. 10.1021/jm051256o.

47. Zhao, H., Koenig, S.G., Dankwardt, J.W., and Singh, S.P. (2014). Practical nonazide synthesis of a D-amino acid oxidase inhibitor via a sequential erlenmeyer-plöchl reaction and ligand-free copper(I) amination protocol. Org Process Res Dev 18, 198–204. 10.1021/op4001737.

48. Ilyin, A.P., Kobak, V. V., Dmitrieva, I.G., Peregudova, Y.N., Kustova, V.A., Mishunina, Y.S., Tkachenko, S.E., and Ivachtchenko, A. V. (2005). Synthesis of annelated azaheterocycles containing a 5-carbamoylpyrazin-3-one fragment by a modification of the four-component Ugi reaction. European J Org Chem, 4670–4679. 10.1002/ejoc.200500522.

