## Supplemental Figures and Tables for "Small-molecule allosteric activator of ubiquitin-specific protease 7 (USP7)"

### Supplemental Figure 1.

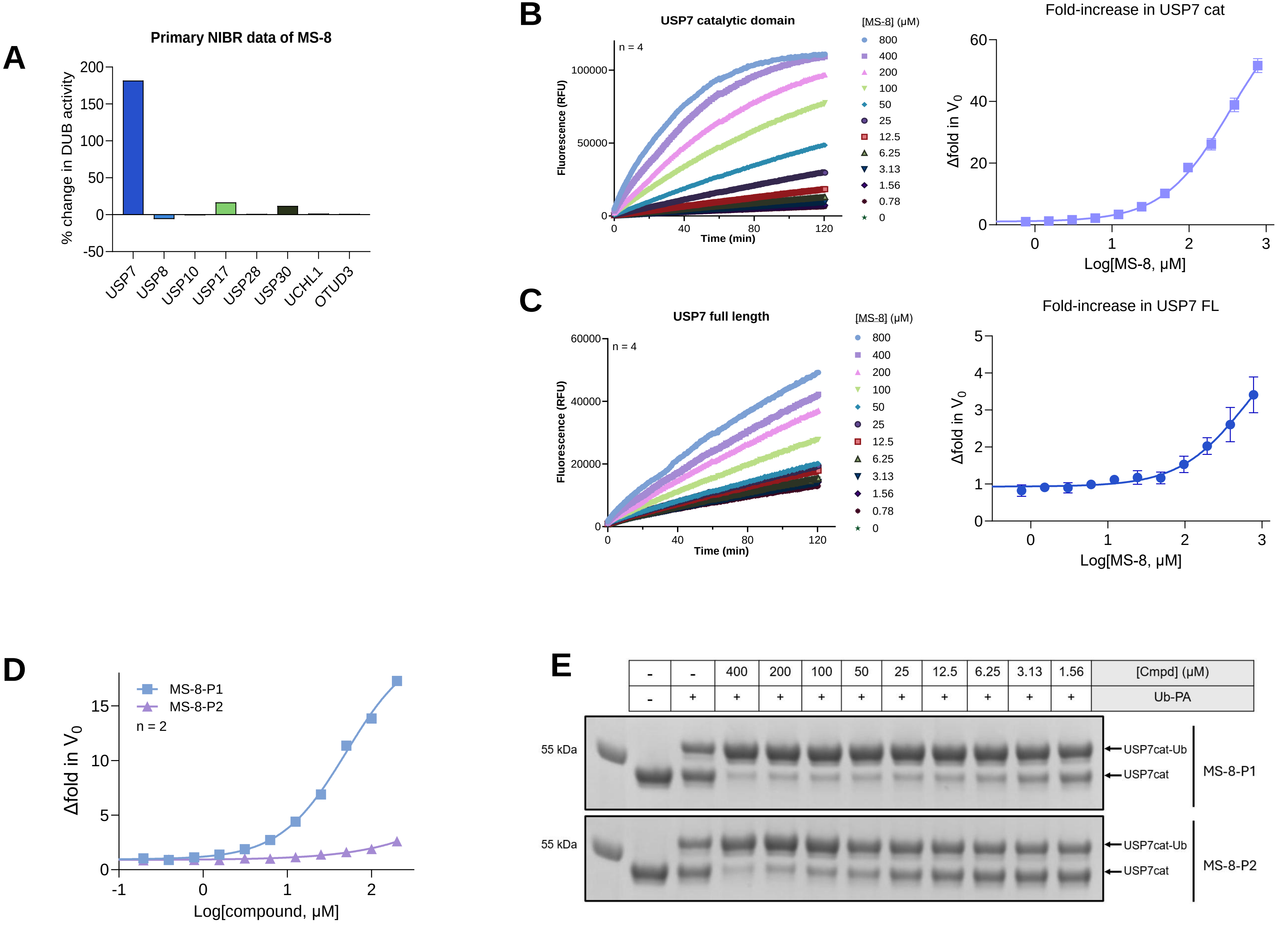

### Supplemental Figure 2.

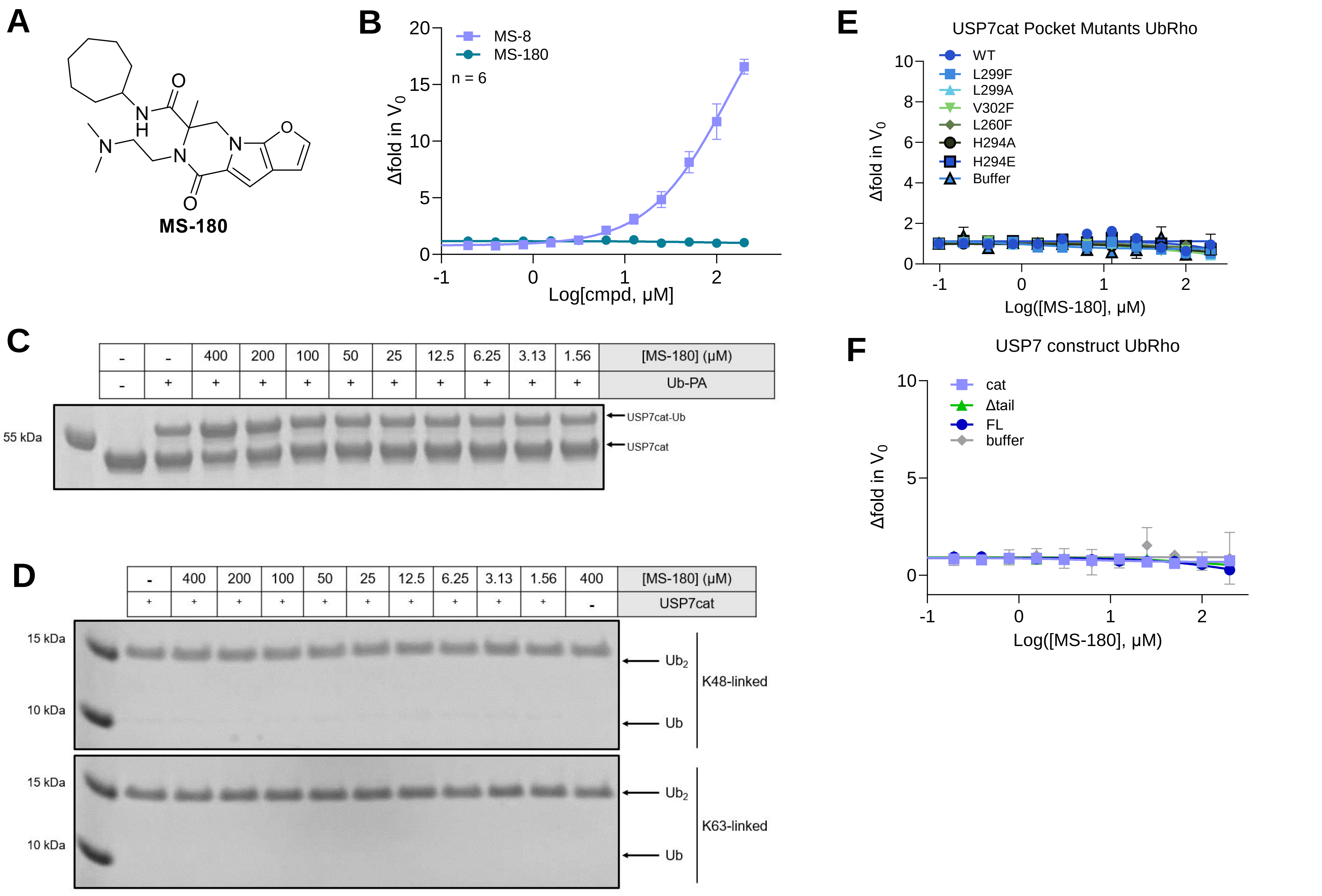

### Supplemental Figure 3.

**A**

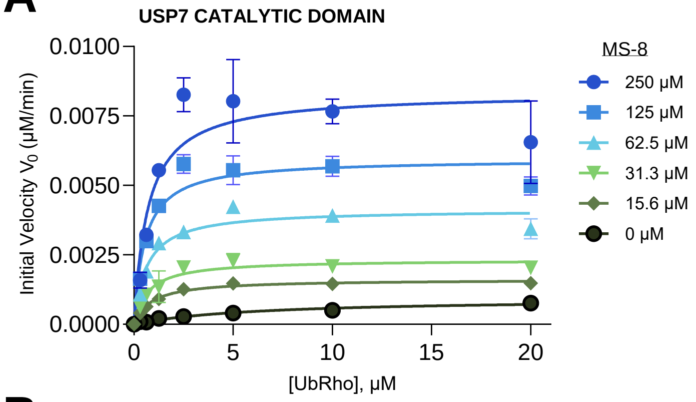


| [MS-8] | 250 μM | 125 μM | 62.5 μM | 31.3 μM | 15.6 μM | 0 μM |
|---|---|---|---|---|---|---|
| $k_{cat}$ ($min^{-1}$) | 0.42 ± 0.04 | 0.30 ± 0.002 | 0.21 ± 0.01 | 0.12 ± 0.01 | 0.08 ± 0.001 | 0.05 ± 0.01 |
| $K_m$ (μM) | 0.70 ± 0.07 | 0.55 ± 0.07 | 0.64 ± 0.01 | 0.73 ± 0.26 | 0.89 ± 0.16 | 6.4 ± 3.18 |

**B**

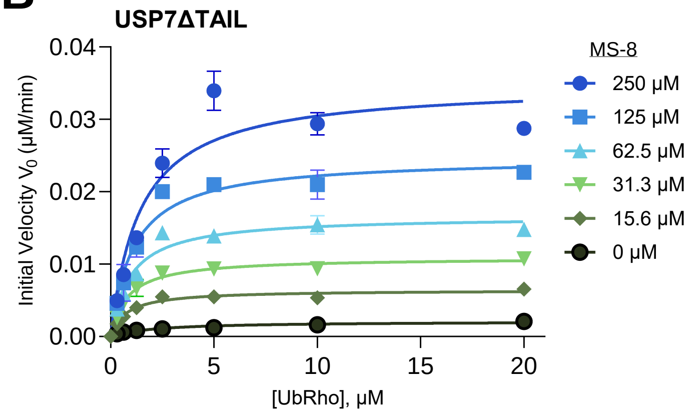


| [MS-8] | 250 μM | 125 μM | 62.5 μM | 31.3 μM | 15.6 μM | 0 μM |
|---|---|---|---|---|---|---|
| $k_{cat}$ ($min^{-1}$) | 1.74 ± 0.09 | 1.26 ± 0.08 | 0.83 ± 0.01 | 0.55 ± 0.01 | 0.32 ± 0.01 | 0.1 ± 0.01 |
| $K_m$ (μM) | 1.44 ± 0.16 | 1.13 ± 0.29 | 0.95 ± 0.06 | 0.83 ± 0.16 | 0.76 ± 0.01 | 2.19 ± 0.4 |

**C**

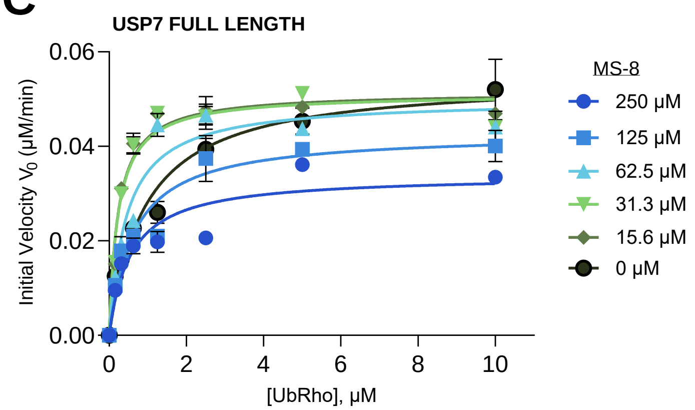


| [MS-8] | 250 μM | 125 μM | 62.5 μM | 31.3 μM | 15.6 μM | 0 μM |
|---|---|---|---|---|---|---|
| $k_{cat}$ ($min^{-1}$) | 3.39 ± 0.09 | 4.28 ± 0.37 | 4.99 ± 0.01 | 5.11 ± 0.13 | 5.15 ± 0.12 | 5.45 ± 0.64 |
| $K_m$ (μM) | 1.1 ± 0.07 | 1.25 ± 0.12 | 0.89 ± 0.01 | 0.46 ± 0.01 | 0.46 ± 0.00 | 1.86 ± 0.28 |

### Supplemental Figure 4.

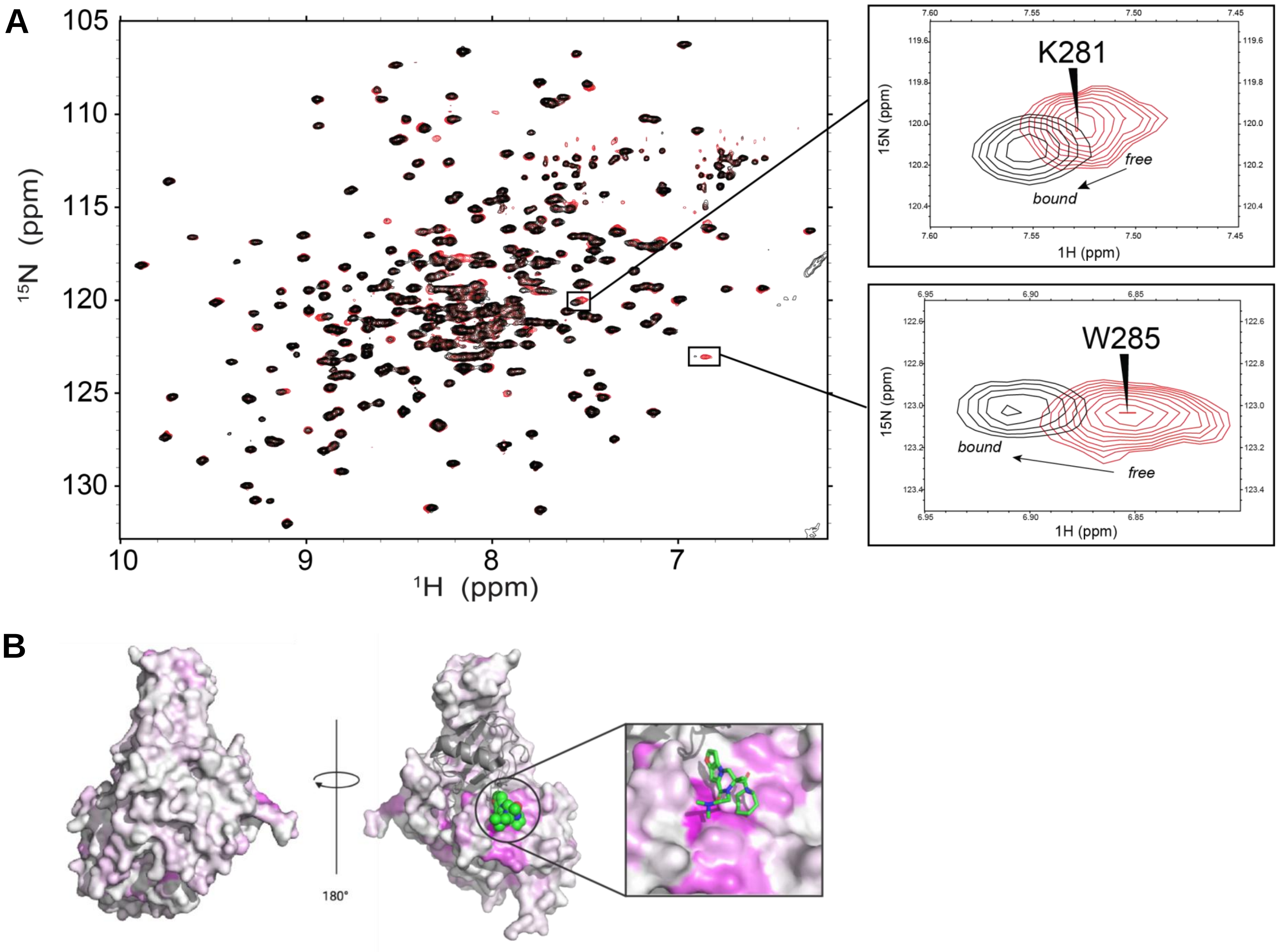

C

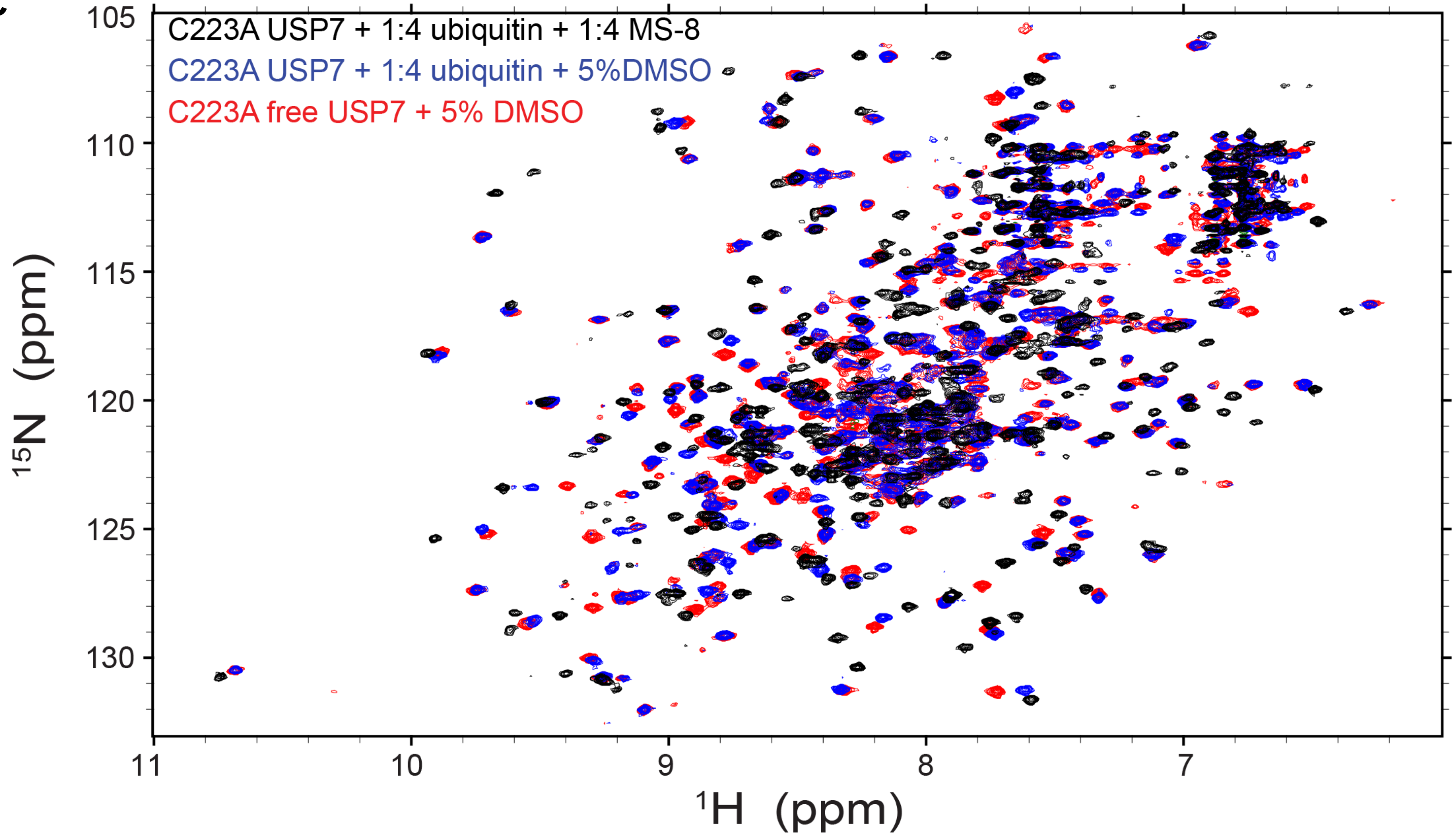
C223A USP7 + 1:4 ubiquitin + 1:4 MS-8
C223A USP7 + 1:4 ubiquitin + 5%DMSO
C223A free USP7 + 5% DMSO
105
110
115
120
125
130
$^{15}$N (ppm)
11
10
9
8
7
$^{1}$H (ppm)

### Supplemental Figure 5.

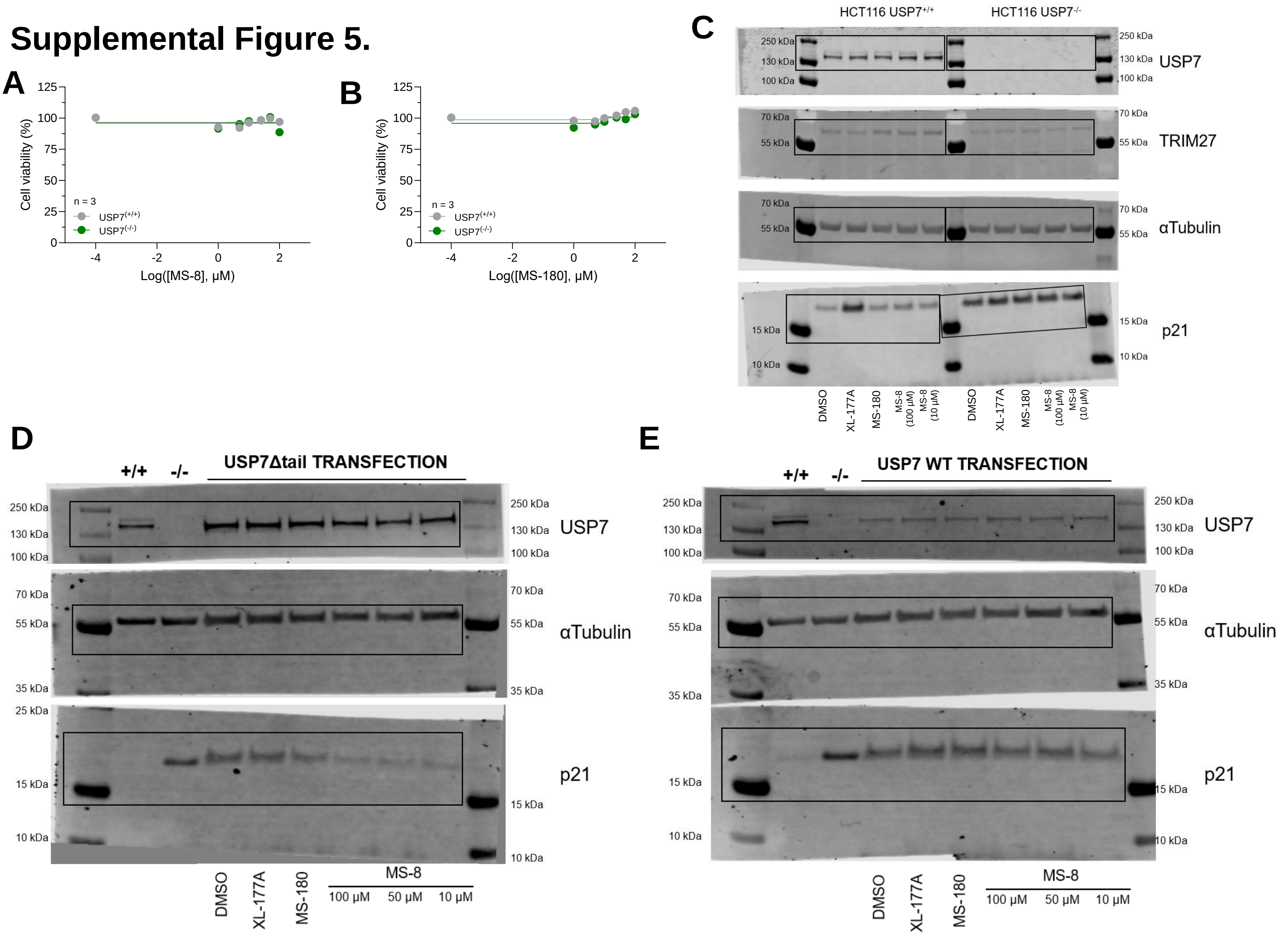

### Supplemental Figure 6.

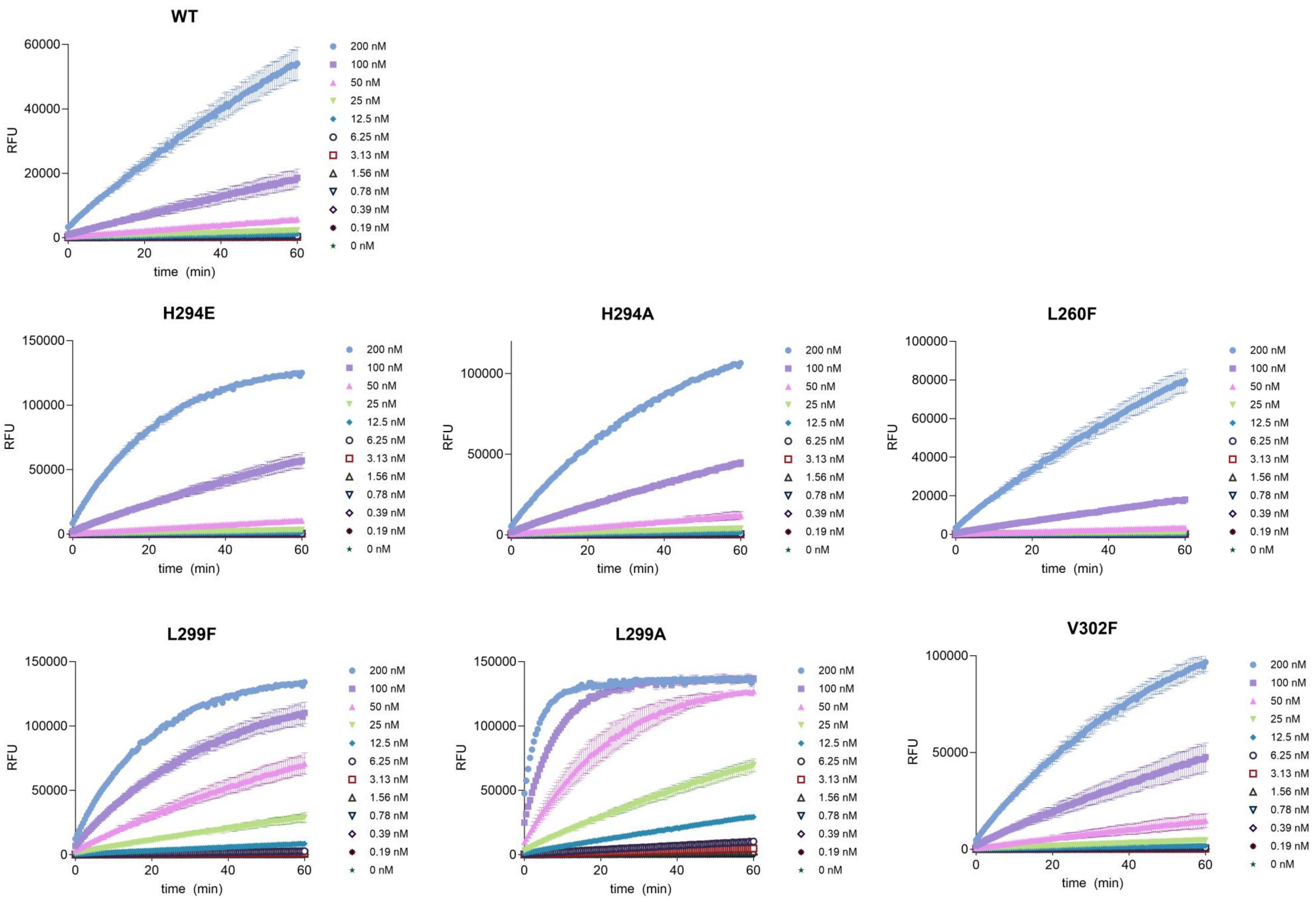

### Supplemental Figure 7.

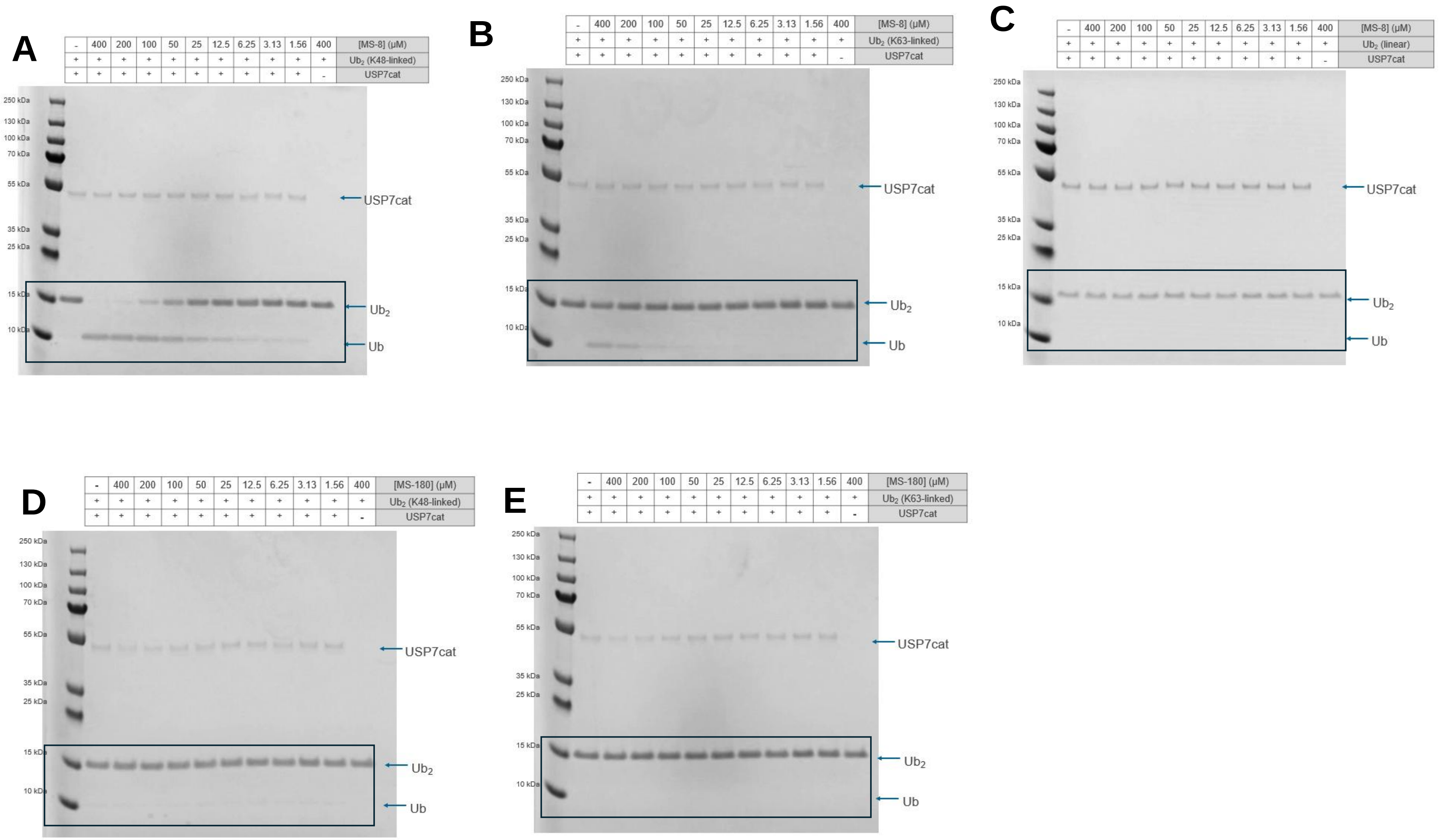

### Supplemental Figure 8.

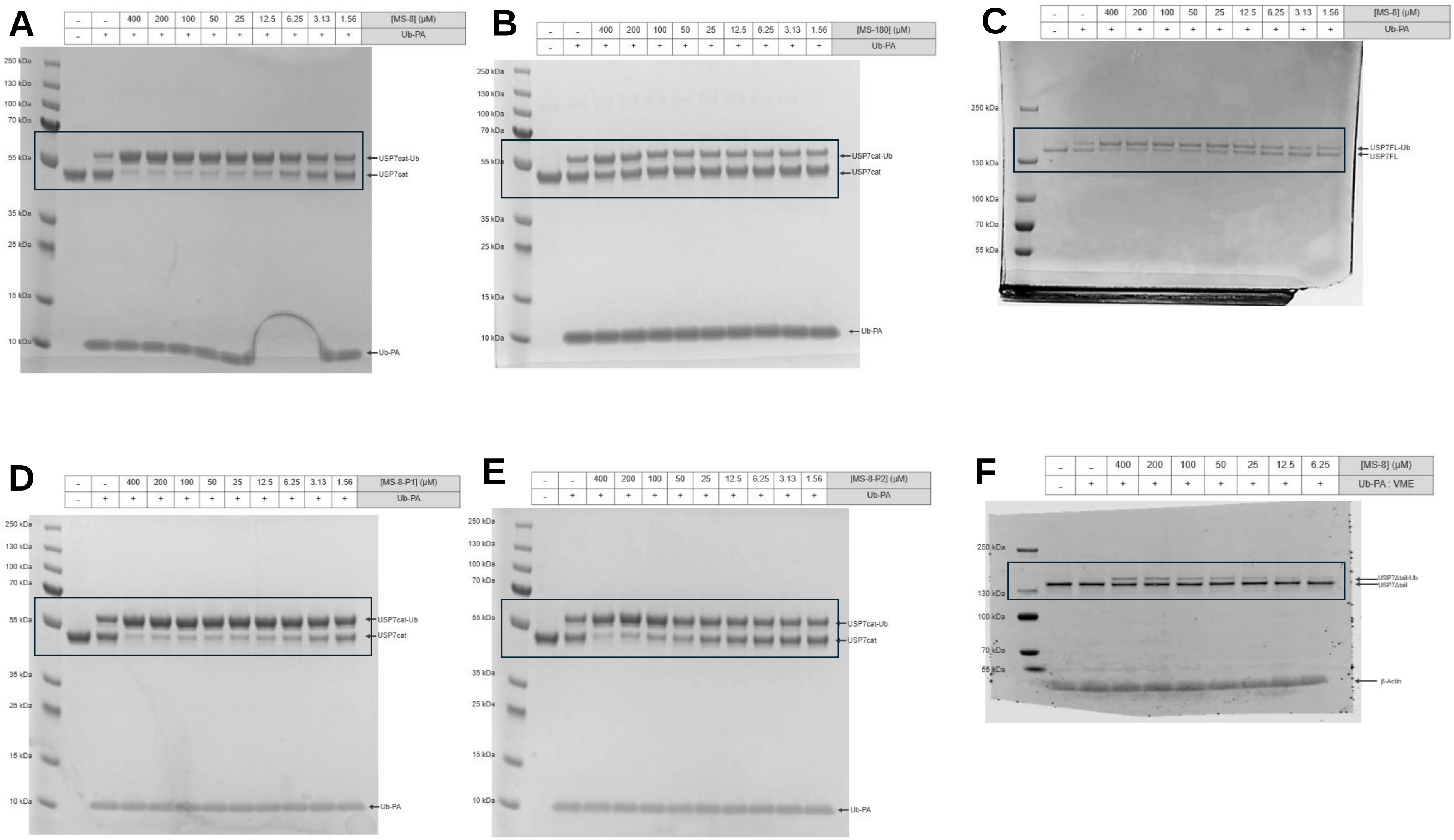

### Supplemental Table 1: Computational supplemental

**A**

| Pocket | Ligand | PDB | box size INNER | box size OUTER | Best docking score (kcal/mol) |
|---|---|---|---|---|---|
| **S3-S5** | MS-8 | 5VS6 | 10 x 10 x 10 | 32 x 32 x32 | -3.90 |
| **Distal** | MS-8 | 5UQX | 10 x 10 x 10 | 26 x 26 x 26 | -5.00 |
| **CPB** | MS-8 | 5UQX | 10 x 10 x 10 | 24 x 24 x 24 | -6.18 |
| **GNE-6776 in distal** | GNE-6776 | 5UQX | 10 x 10 x 10 | 26 x 26 x 26 | -8.10 |

**B**

| Pocket | Ligand | atoms | Simulation Box (Å Å Å) | Simulation Length (ns) |
|---|---|---|---|---|
| **S3-S5** | MS-8 | 52272 | 95 X 74 X 67 | 100 X 3 |
| **Distal** | MS-8 | 48392 | 94 X 73 X 69 | 100 X3 |
| **CPB** | MS-8 | 49429 | 97 X 80 X 66 | 100 X 3 |
| **GNE-6776 in distal** | GNE-6776 | 51140 | 97 X 72 X 69 | 100 X 1 |

### Supplemental Table 2. Mutagenesis primers

**A**

| USP7 CAT Mutations | Forward Primer Sequence | Reverse Primer Sequence |
|---|---|---|
| H294E | CTTCATGCAAGAAGATGTTCAGG | CTATCTAAAGTTTCCCACC |
| H294A | CTTCATGCAAGCGGATGTTCAGGAGCTTTG | CTATCTAAAGTTTCCCACC |
| L260F | CCCTTTAGCATTTCAAAGAGTGTTC | ACGCTTTTAGACGAATC |
| L299F | TGTTCAGGAGTTTTGTCGAGTGT | TCATGTTGCATGAAGCTATC |
| L299A | GTTCAGGAGGCGTGTCGA | TCGACACGCCTCCTGAAC |
| V302F | GCTTTGTCGAtttTTGCTCGATAATG | TCCTGAACATCATGTTG |

**B**

| USP7 FL Mutations | Forward Primer Sequence | Reverse Primer Sequence |
|---|---|---|
| ΔTAIL (aa 1 – 1083) | TGATCTAGAGTCGACCCG | GTTGAAGTGGTCGAGCCC |

**C**

| USP7 | Treatment | $K_D$ (μM) |
|---|---|---|
| Full length | DMSO | 55.48 |
| Catalytic domain | DMSO | 198.8 |
| | 25 μM MS-8 | 148.5 |
| | 100 μM MS-8 | 132.9 |
